# Predictive regression modeling with MEG/EEG: from source power to signals and cognitive states

**DOI:** 10.1101/845016

**Authors:** David Sabbagh, Pierre Ablin, Gaël Varoquaux, Alexandre Gramfort, Denis A. Engemann

## Abstract

Predicting biomedical outcomes from Magnetoencephalography and Electroencephalography (M/EEG) is central to applications like decoding, brain-computer-interfaces (BCI) or biomarker development and is facilitated by supervised machine learning. Yet most of the literature is concerned with classification of outcomes defined at the event-level. Here, we focus on predicting continuous outcomes from M/EEG signal defined at the subject-level, and analyze about 600 MEG recordings from Cam-CAN dataset and about 1000 EEG recordings from TUH dataset. Considering different generative mechanisms for M/EEG signals and the biomedical outcome, we propose statistically-consistent predictive models that avoid source-reconstruction based on the covariance as representation. Our mathematical analysis and ground truth simulations demonstrated that consistent function approximation can be obtained with supervised spatial filtering or by embedding with Riemannian geometry. Additional simulations revealed that Riemannian methods were more robust to model violations, in particular geometric distortions induced by individual anatomy. To estimate the relative contribution of brain dynamics and anatomy to prediction performance, we propose a novel model inspection procedure based on biophysical forward modeling. Applied to prediction of outcomes at the subject-level, the analysis revealed that the Riemannian model better exploited anatomical information while sensitivity to brain dynamics was similar across methods. We then probed the robustness of the models across different data cleaning options. Environmental denoising was globally important but Riemannian models were strikingly robust and continued performing well even without preprocessing. Our results suggest each method has its niche: supervised spatial filtering is practical for event-level prediction while the Riemannian model may enable simple end-to-end learning.

## 1. Introduction

Magnetoencephalography and Electroencephalography (M/EEG) access population-level neuronal dynamics across multiple temporal scales from seconds to milliseconds (Buzsáki and Draguhn, 2004; Hämäläinen et al., 1993). Its wide coverage of brain rhythms supports modeling cognition and brain health at different levels of organization from states to traits (Baillet, 2017; Buzsáki and Watson, 2012; da Silva, 2013). In the past decades, this has led to predictive modeling approaches in which cognitive or clinical outcomes are statistically approximated from the electrophysiological signals (Besserve et al., 2007; Woo et al., 2017). In a common scenario, single-trial stimulus details are predicted from chunks of event-level signal, *e.g.*, visual orientation or auditory novelty (Cichy et al., 2015; King et al., 2013). With brain-computer-interfaces (BCI), one aims to read out cognitive states and translate them into control signals, e.g., to capture movement-intentions (Lotte et al., 2007; Tangermann et al., 2008; Wolpaw et al., 1991). For biomarkers applications, the focus is on predicting medical diagnosis and other clinical endpoints (Engemann et al., 2018; Mazaheri et al., 2018; Sami et al., 2018).

What is the physiological source of M/EEG-based prediction? Similar to an analog radio, M/EEG receives signals containing multiplexed streams of information in different frequency ‘channels’ (Akam and Kullmann, 2014; Panzeri et al., 2010; van Wassenhove, 2016). The signal comprises periodic and arrhythmic components which give rise to the characteristic 1/f power law regime (Dehghani et al., 2010; He et al., 2010; Linkenkaer-Hansen et al., 2001). M/EEG brain-dynamics originate from transient large-scale synchrony of distinct brain-networks where the anatomical regions involved communicate in different frequency bands (Hipp et al., 2012; Siegel et al., 2012). Typically, the frequency depends on the spatial scale of the network: as the scale becomes more local the spectral frequency increases (Buzsáki and Draguhn, 2004; Honey et al., 2007).

This motivates modeling approaches sensitive to both the temporal scale and the topography of the signal. Unfortunately, the neural sources of M/EEG cannot be observed and have to be inferred with uncertainty from their distorted representation on extra-cranial sensors. This argues in favor of statistical-learning techniques that can readily exploit high-density sensor arrays beyond sensor-wise statistical testing. So far this has been approached by explicit biophysical source modeling (Hämäläinen and Ilmoniemi, 1994; Khan et al., 2018; Westner et al., 2018), statistical approximations of biophysical generators through Independent Component Analysis (ICA) (Hyvärinen and Oja, 2000; Makeig et al., 1995; Stewart et al., 2014; Subasi and Gursoy, 2010), spatial filtering approaches often inspired by BCI (Dähne et al., 2013, 2014a,b; de Cheveigné and Parra, 2014; Haufe et al., 2014a; Nikulin et al., 2011) or direct application of general purpose machine learning on the sensor time series (King et al., 2013).

Strikingly, the bulk of the literature on predictive modeling from M/EEG focuses on classification problems, evoked responses and outcomes defined at the event-level, e.g., brain responses to stimuli or brain dynamics related motor behavior. This is understandable, as evoked response analysis draws on rich resources from a long-standing history in experimental psychology (Coles and Rugg, 1995; Näätänen, 1975; Polich and Kok, 1995) and lend themselves to categorical problems as defined by experimental conditions. Besides, working on classification rather than regression may be more rewarding, as learning the boundary between classes is easier than estimating a full regression function (Hastie et al., 2005, chapter 7.3.1). Nevertheless, high-interest clinical outcomes other than diagnosis are often continuous and often involve predicting at the subject-level (e.g., prediction of risk scores, optimal drug-dosage, time of hospitalization or survival). Moreover, as EEG-recordings are combined across medical sites where different EEG-protocols are used, additional strain is put on spontaneous brain rhythms that can be accessed even if no particular task is used (Engemann et al., 2018). Yet, it is currently unclear how learning approaches based on brain rhythms compare as the data generating mechanism changes (event-level vs subject-level outcomes) or when the underlying probability model (*e.g.* log-linear vs linear relationship to power) is not a priori known.

In this paper we focus on linking neural power spectra with their measure in M/EEG using appropriate models that facilitate prediction with high-dimensional regression. We aim to answer the following questions: 1) How can regression on M/EEG power-spectra be related to statistical models of the outcome and the neural signal? 2) What are the mathematical guarantees that a type of regression captures a given brain-behavior link? 3) How do ensuing candidate models perform in the light of model violations, uncertainty about the true data-generating process, variable noise, and different preprocessing options? The article is organized as follows: First we detail commonly used approaches for M/EEG-based predictive modeling. Subsequently, we develop a coherent mathematical framework for relating M/EEG-based regression to models of the neural signal, and, as a result, propose to conceptualize regression as predicting from linear combinations of uncorrelated statistical sources. Then we present numerical simulations which confront different regression models with commonly encountered model violations. Subsequently, we conduct detailed model comparisons on MEG and EEG data for event-level and subject-level problems. Finally, we investigate practical issues related to availability of source-modeling and preprocessing options.

## 2. Methods

### 2.1. State-of-the art approaches to predict from M/EEG observations

One important family of approaches for predictive modeling with M/EEG is relying on explicit biophysical source modeling. Here, anatomically constrained inverse methods are used to infer the most likely electromagnetic source configuration given the observations (Hämäläinen et al., 1993). Common techniques rely on fitting electrical-current dipoles (Mosher et al., 1992) or involve penalized linear inverse models to estimate the current distribution over a pre-specified dipole grid (Hämäläinen and Ilmoniemi, 1994; Hauk and Stenroos, 2014; Lin et al., 2006; Van Veen and Buckley, 1988). Anatomical prior knowledge is injected through the well-defined forward model: Maxwell equations enable computing leadfields from the geometry and composition of the head, which predict propagation from a known source to the sensors (Hämäläinen et al., 1993; Mosher et al., 1999). From a signal-processing standpoint, when these steps lead to a linear estimation of the sources, they can be thought of as biophysical spatial filtering. Prediction is then based on the estimated source-signals, see for example (Khan et al., 2018; Kietzmann et al., 2019; Westner et al., 2018).

A second family is motivated by unsupervised decomposition techniques such as Independent Component Analysis (Hyvärinen and Oja, 2000; Makeig et al., 1996), which also yield spatial filters and estimates of maximally independent sources that can be used for prediction (Stewart et al., 2014; Subasi and Gursoy, 2010; Wang and Makeig, 2009). Such methods model the data as an independent set of statistical sources that are entangled by a so-called mixing matrix, often interpreted as the leadfields. Here, the sources are purely statistical objects and no anatomical notion applies directly. In practice, unsupervised spatial filters are often combined with source modeling and capture a wide array of situations ranging from single dipole-sources to entire brain-networks (Brookes et al., 2011; Delorme et al., 2012; Hild II and Nagarajan, 2009).

Finally, a third family directly applies general-purpose machine learning on sensor space signals without explicitly considering the data generating mechanism. Following a common trend in other areas of neuroimaging research (Dadi et al., 2019; He et al., 2019; Schulz et al., 2019), linear prediction methods have turned out extraordinarily well-suited for this task, *i.e.*, logistic regression (Andersen et al., 2015), linear discriminant analysis (Wardle et al., 2016), linear support vector machines (King et al., 2013).

The success of linear models deserves separate attention as these methods enable remarkable predictive performance with simplified fast computation (Parra et al., 2005). While interpretation and incorporation of prior knowledge remain challenging, significant advances have been made in the past years. This has led to novel methods for specifying and interpreting linear models (Haufe et al., 2014b; van Vliet and Salmelin, 2019). Recent work has even suggested that for the case of learning from evoked responses, linear methods are compatible with the statistical models implied by source localization and unsupervised spatial filtering (King and Dehaene, 2014; King et al., 2018; Stokes et al., 2015). Indeed, if the outcome is linear in the source signal, *i.e.*, due to the linear superposition principle, the mixing amounts to a linear transform that can be captured by a linear model with sufficient data. Additional source localization or spatial filtering should therefore be unnecessary in this case.

On the other hand, the situation is more complex when predicting outcomes from brain rhythms, *e.g.*, induced responses (Tallon-Baudry and Bertrand, 1999) or spontaneous oscillations. As brain-rhythms are not strictly time-locked to external events, they cannot be accessed by averaging. Instead, they are commonly represented by the signal power in shorter or longer time windows and often give rise to log-linear models (Buzsáki and Mizuseki, 2014; Roberts et al., 2015). A consequence of such non-linearities is that it cannot be readily captured by a linear model. Moreover, simple strategies such as log-transforming the power estimates only address the issue when applied at the source-level: the leadfields have already spatially smeared the signal presented on the sensors.

This leads back to spatial filtering approaches. Beyond source localization and unsupervised filtering, supervised spatial filtering methods have recently become more popular beyond the context of BCIs. These methods solve generalized eigenvalue problems to estimate coordinate systems constructed with regard to criteria relevant for prediction. For example, spatio-spectral-decomposition (SSD) is an unsupervised technique that enhances SNR with regard to power in surrounding frequencies (Nikulin et al., 2011). On the other hand, Common Spatial Patterns (Koles, 1991) and Joint Decorrelation (de Cheveigné and Parra, 2014) or Source Power Comodulation (SPoC) focus on correlation with the outcome (Blankertz et al., 2008; Dähne et al., 2013, 2014a), whereas Dmochowski et al. (2012) have proposed variants of Canonical Correlation Analysis (CCA) (Dähne et al., 2014b; Hotelling, 1992) without orthogonality constraint to focus on shared directions of variation between related datasets or by proposing shared envelope correlations as optimization target (Dähne et al., 2014b). This yields a two-step procedure: 1) spatial filters model the correlation induced by the leadfields and provide unmixed time-series 2) some non-linear transforms such as logarithms are applied to these time-series as the validity of linear equations is now secured.

A more recent single-step approach consists in learning directly from spatially correlated power-spectra with linear models and Riemannian geometry (Barachant et al., 2011, 2013; Fruehwirt et al., 2017; Rodrigues et al., 2019; Yger et al., 2017). This mathematical framework provides principles to correct for the geometric distortions arising from linear mixing of non-linear sources. This is achieved by using a Riemannian metric, immune to linear transformations, which allows representing the covariance matrices used for representing the M/EEG signal as Euclidean objects for which linear models apply. This approach has turned out to be promising for enhancing classification of event-level data and has been the important ingredient of several winning solutions in recent data analysis competitions, *e.g.*, the seizure prediction challenge organized by the University of Melbourne in 2016. Recently, this approach has been explored for prediction of subject-level brain volume from clinical EEG in Alzheimer’s disease in about 100 patients (Fruehwirt et al., 2017). Yet, systematic comparisons against additional baselines and competing regression models on larger datasets and other outcomes are missing.

Importantly, the majority of approaches have focused on event-level prediction problems instead of subject-level prediction and have never been been systematically compared in terms of their statistical properties and empirical behavior. Here we will explicitly focus on subject-level as contrasted to event-level prediction, both, theoretically and at the level of data analysis. Note that the present article does not focus on event-level prediction with generalization across subjects (Halme and Parkkonen, 2018; Olivetti et al., 2014; Westner et al., 2018), which is a distinct and more complex problem inheriting its structure from, both, event-level and subject-level regression. A summary of mathematical notations used in this article can be found in table 1.

**Table 1:**
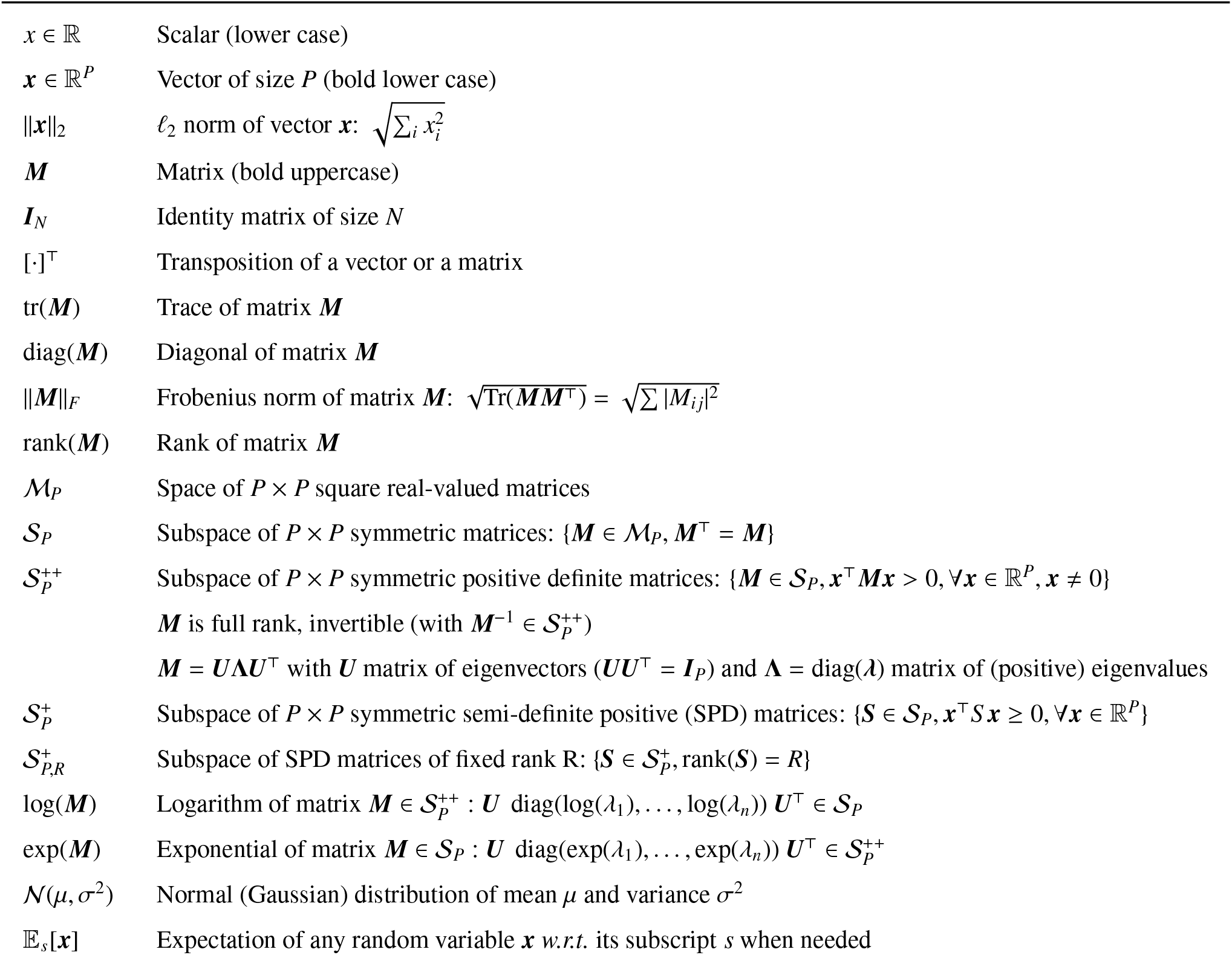
Mathematical notations used in this article

### 2.2. A priori knowledge: the biophysical data-generating mechanism

MEG and EEG signals are produced by electrical sources in the brain that emerge from the synchronous activity of cortical layer IV pyramidal neurons (Hämäläinen et al., 1993). These neural current generators form *physiological sources* in the brain. We will assume the existence of *M* such sources, *z*(*t*) ∈ ℝ^*M*^, where *t* represents time. These sources can be thought of as localized current sources, such as a patch of cortex with synchronously firing neurons, or a large set of patches forming a network. In this work, we are interested in predicting an outcome *y* ∈ ℝ from multivariate MEG/EEG signals ***x***(*t*) ∈ ℝ^*P*^, where *P* corresponds to the number of sensors. The underlying assumption is that the physiological sources are at the origin of the signals ***x***, and that they are statistically related to *y*. Often they are even the actual generators of *y*, *e.g.*, the neurons producing the finger movement of a person. Here, we embrace the statistical machine learning paradigm where one aims to learn a predictive model from a set of *N* labeled training samples, (***x***_*i*_(*t*), *y_i_*), *i* = 1,…, *N*, which we see, fundamentally, as a function approximation problem. We will consider predicted outcomes that do not dependent on time. The physics of the problem and the linearity of the quasi-static approximation of Maxwell’s equations guarantee that MEG/EEG acquisition is linear too: the signals measured are obtained by linear combination of the underlying physiological sources. This leads to:

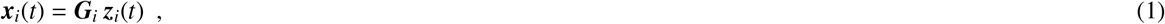

where ***G***_*i*_ ∈ ℝ^*P×M*^ is the leadfield, also commonly referred to as gain matrix. Therefore, the observed M/EEG signal ***x**_i_*(*t*) contains information on unobserved brain sources ***z**_i_*(*t*), distorted by individual brain anatomy represented by ***G**_i_*. Note that here the *j*-th column of ***G**_i_* is not necessarily constrained to be the forward model of a focal electrical current dipole in the brain. It can also correspond to large distributed sources. This reality is illustrated as the area outside the cloud in Fig. 1.

**Figure 1:**
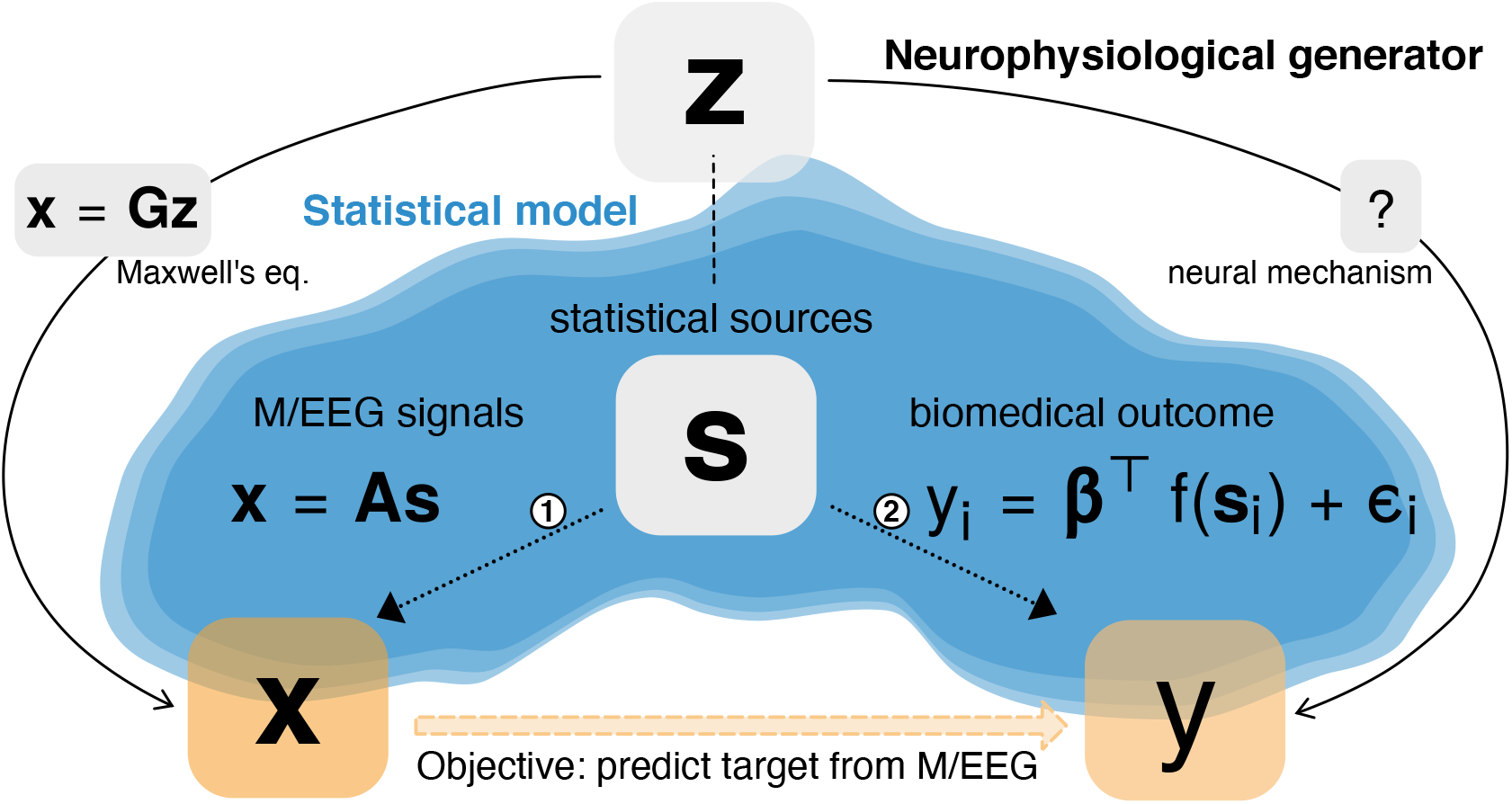
Generative model for regression with M/EEG. Unobservable neuronal activity *z* gives rise to observed M/EEG data ***X*** and an observed biomedical outcome *y*. The M/EEG data **X** is obtained by linear mixing of *z* through the leadfield ***G***. The outcome *y* is derived from *z* through often unknown neural mechanisms. The statistical model (blue cloud) approximates the neurophysiological data-generating mechanisms with two sub-models, one for the M/EEG signals ***X*** (path 1), one for the biomedical outcome *y* (path 2). Both models are based on a vector *s* of uncorrelated statistical sources that, may refer to localized cortical activity or synchronous brain networks. The ensuing model generates *y* from a linear combination of the statistical sources *s*. The generative model of **X** follows the ICA model (Hyvärinen and Oja, 2000) and assumes linear mixing of the source signals by ***A***, interpreted as a linear combination of the columns of the leadfield **G**. The generative model of *y* assumes a linear model in the parameters *β* but allows for non-linear functions in the data, such as the power or the log-power. The mechanisms governing path 1 implies that the sources s appear geometrically distorted in **X**. This makes it impossible for a linear model to capture this distortion if *y*, in path 2, is generated by a non-linear function of s. This article focuses on how to mitigate this distortion without biophysical source modeling when performing regression on M/EEG source power.

A natural approach to estimate a regression model in this setting consists in estimating the locations, amplitudes and extents of the sources from the MEG/EEG data. This estimation known as in the inverse problem (Baillet, 2017) can be achieved using Minimum Norm Estimates (MNE) (Hämäläinen and Ilmoniemi, 1984). From the estimated sources, one can then learn to predict y as the distortions induced by individual head geometry are mitigated. While approaching the problem from this perspective has important benefits, such as the ability to exploit the geometry and the physical properties of the head tissues of each subject, there are certain drawbacks. First, the inverse problem is ill-posed and notoriously hard to solve. Second, computing ***G**_i_* requires costly MRI acquisitions and time-consuming manual labor by experienced MEG/EEG practitioners: anatomical coregistration and tedious data-cleaning to mitigate electromagnetic artefacts caused by environmental or physiological sources of non-interest outside of the brain. The purpose of this article is to show how to learn a regression model without biophysical source modeling. Note that, in this article, we use the term *generative model* in the statistical sense of a probabilistic model of the M/EEG observations and the biomedical outcomes.

### 2.3. Generative models: statistical approximation of the outcome and the M/EEG signals

Independent Component Analysis (Hyvärinen and Oja, 2000, ICA) is another popular approach to model M/EEG signals (Makeig et al., 1997). We consider *Q* ≤ *P statistical sources **s***(*t*) ∈ ℝ^*Q*^, that correspond to unknown latent variables. These variables are assumed to be statistically related to the outcome variable *y* and to be linearly related to measured signal **x**(*t*). The area inside the cloud depicted in Fig. 1 illustrates the generative models.

#### Generative model of the M/EEG signals

We consider an extension of noise-free Blind Source Separation (Belouchrani et al., 1997). We assume the signal arises from activity of uncorrelated sources, contaminated by an additive noise. The sources span the same space for all samples, of dimension *Q*. This model is conveniently written in matrix form:

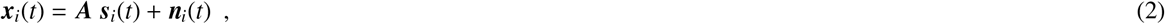

where ***s**_i_*(*t*) ∈ ∝^*Q*^ is the time-series of sources amplitude of sample *i* and ***n**_i_*() ∈ ∝^*P*^ is the contamination due to noise. The columns of the *mixing matrix **A*** ∈ ∝^*P×Q*^ are the *Q* linearly independent *source patterns* (Haufe et al., 2014b), which correspond to topographies on the sensor array. Each quantity in the right-hand side of Eq. (2), **A**, ***s**_i_*(*t*) and ***n**_i_*(), is unknown and should be inferred from ***x**_i_*(*t*). This setting encompasses both event-level regression, where the samples ***x**_i_*(*t*) are epochs of signal from a unique subject (i stands for a particular time window), and subject-level regression where the samples represent the full signal of multiple subjects (*i* then stands for a particular subject). In this latter case, the assumption that ***A*** is fixed across samples is not realistic but useful for modeling purposes. Model violations will be addressed in section 2.5.

We also make the assumption that the noise subspace is not “mixed” with the source subspace and that the noise subspace is shared across samples. This is motivated by the fact that environmental events generate the strongest noise in M/EEG recordings. These environmental perturbations are by definition independent from brain activity. On the other hand, physiological noise, due to cardiac or ocular activity, systematically interacts with brain signals and is necessarily captured by the statistical sources ***s***. We can then extend ***A*** to include source and noise patterns, making it an invertible matrix ***A*** ∈ ℝ^*P×P*^. Denoting ***η**_i_*(*t*) ∈ ℝ^*P*^ the concatenation of source and noise signals, the generative model (2) can be compactly rewritten as:

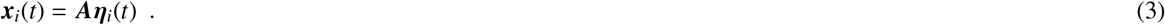

Justification of this formulation can be found in Appendix 5.1. Note that this statistical generative model is a simplification of the biophysical generative mechanism: the real sources ***z**_i_* may not be independent (Nolte et al., 2006), the gain ***G**_i_* is sample-dependent in subject-level regression, and the number of true sources may exceed the number of sensors, *M* >> *P*.

#### Generative model of the biomedical outcome

The proposed framework models *y_i_* as a function of the sources powers:

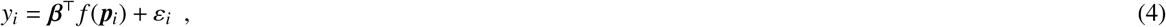

where 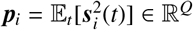 is the power of sources of sample *i, f* is a known function, ***β*** ∈ ℝ^*Q*^ are regression coefficients, and *ε_i_* is an additive random perturbation. In practice, one prefers frequency specific models, where the previous relationships are obtained after ***s**_i_*(*t*) has been bandpass filtered in a specific frequency range. The broadband covariance (computed on the raw signal without temporal filtering) largely reflects low-frequency power as consequence of the predominant 1/f power spectrum, hence, is rarely of interest for predicting. In frequency-specific models, the powers are replaced by band-powers: power of the source in the chosen frequency band. Note that source power in a given frequency band is simply the variance of the signal in that frequency band. The noise in the outcome variable depends on the context: it can represent intrinsic measurement uncertainty of *y_i_*, for example sampling rate and latency jitter in behavioral recordings, inter-rater variability for a psychometric score, or simply a model mismatch. The true model may not be linear for example.

Linear models in the sources powers (*f* = identity) or log-powers (*f* = log) are commonly used in the literature and support numerous statistical learning models on M/EEG (Blankertz et al., 2008; Dähne et al., 2014a; Grosse-Wentrup* and Buss, 2008). In particular Buzsáki and Mizuseki (2014) discusses a wide body of evidence arguing in favor of log-linear relationships between brain dynamics and cognition. Both possibilities for *f* will be considered below.

### 2.4. Regression models

In this article, we use a statistical learning approach and focus on the function approximation problem rather than parameter estimation. We aim at finding regression models that are *statistically consistent i.e.* with no approximation error. Any regression model has a generalization error that essentially entails three terms: the approximation error (when the true function linking *x* and *y* does not belong to the search space of our regression model), the estimation error (when learning from a finite random sample), the irreducible error (when the relation between *y* and *x* is not deterministic). A statistically consistent regression model is a model with no approximation error. When learning from a sufficiently large number of samples (no estimation error) and without noise in *y* (no irreducible error), a regression model with no approximation error will have no generalization error. It has then learnt a function that perfectly approximates the true function asymptotically.

The generative model (4) of the outcome involves source powers *i.e.* the squared amplitude of the source signal. This non-linear dependence hints at using non-linear features of the M/EEG signal ***x***_*i*_, based on its second-order moment: the between-sensors covariance matrix. Denoting the data matrix ***X**_i_* ∈ ℝ^*P×T*^ with *T* the number of time samples, the covariance matrix of signal ***x_i_***(*t*) (assumed to be zero-mean) reads:

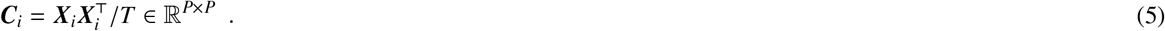

The diagonal of this matrix represents the variance (power) of each sensor, while the off-diagonal terms contain the covariance between each pair of signals. Negative values in the off-diagonal express negative correlations. If we assume that the components of the sources are zero-mean and uncorrelated, the covariance matrix of sources is diagonal: 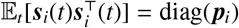. If they are also uncorrelated from the noise 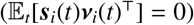, then the covariances have a specific structure given by:

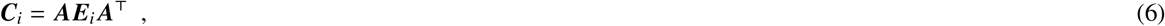

where 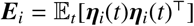 is a block diagonal matrix, whose upper *Q* × *Q* block is the covariance of sources diag(***p**_i_*) and the (*P - Q*) × (*P - Q*) lower block is the covariance of the noise. In particular, these covariances ***C**_i_* are full rank. They contain information about source powers ***p**_i_* in ***E**_i_* but this information is noisy and distorted through unknown linear field spread **A**. This unknown mixing makes it challenging to find optimal regression models with no approximation error.

In this work, we introduce three different regression models that successfully achieve statistical consistency for certain generative assumptions. They are all based on a linear model, applied to carefully chosen non-linear feature vector ***v**_i_*, based on the covariance ***C**_i_*.

#### Upper regression model

This model consists in taking

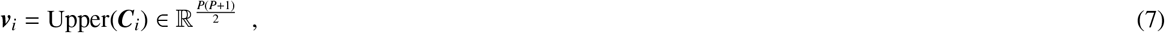

where Upper(***M***) is defined as the vector containing the upper triangular coefficients of ***M***, with off-diagonal terms weighted by a factor 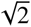. This weighting ensures that the vector and the matrix have same norms (||Upper(***M***)||_2_ = ||***M***||_*F*_). The ‘upper’ model is consistent in the particular case where *f* = identity. Indeed, rewriting Eq. (6) as ***E_i_*** = ***A***^-1^***C**_i_**A***^-τ^, and since the *p_i,j_* are on the diagonal of the upper block of ***E**_i_*, the relationship between the ***p**_i,j_* and the coefficients of ***C**_i_* is also linear. Since the variable of interest *y_i_* is linear in the coefficients of ***p**_i_*, it is also linear in the coefficients of ***C**_i_*, hence linear in the coefficients of ***v**_i_*. In other words, the ‘upper’ regression model is statistically consistent for *f* = identity. This method cannot be generalized to an arbitrary spectral function *f* because *f*(**C**_*i*_) ≠ **A** *f*(**E_*i*_**) **A**^*τ*^.

#### Riemann regression model

The Riemannian embedding yields a representation of sensor-level power and its correlation structure relative to a common reference. In the particular case where *f* = log, the idea is to normalize each covariance ***C**_i_* by a common reference 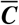, the geometric mean of covariances ***C**_i_*. Then we can show that a linear model applied to feature vector:

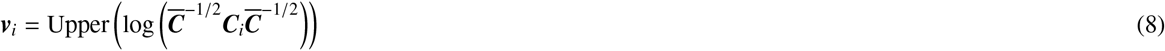

leads to a consistent regression model (see proof in Appendix 5.3). This, essentially, means taking the log of ***C**_i_* after it has been whitened by 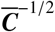, making the quantity of interest relative to some reference 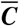. This formulation has also a nice interpretation in terms of Riemannian geometry as being the projection of covariance matrix ***C**_i_* to a common Euclidean space: the tangent space at 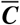. A concise and self-contained introduction to Riemannian manifolds can be found in Appendices 5.4 and 5.5. In particular the norm of ***v**_i_* can be interpreted as the (geometric) distance between ***C**_i_* and 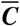 and does not depend on **A**. Essentially, the Riemannian approach projects out fixed linear spatial mixing through the whitening with the common reference. Finally, even though the geometric mean is the most natural reference on the positive definite manifold, consistency of the Riemann regression model still holds when using the Euclidean mean as the common reference point. Indeed, a recent study on fMRI-based predictive modeling has reported negligible differences between the two options (Dadi et al., 2019, appendix A).

#### SPoC regression model

The *SPoC* regression model achieves consistency for any function *f* by taking a rather different approach: it consists in recovering the inverse of the mixing matrix ***A***. The SPoC algorithm is a supervised spatial filtering algorithm simultaneously discovered by de Cheveigné and Parra (2014) and Dähne et al. (2014a). In general, spatial filtering consists in computing linear combinations of the signals to produce so-called *virtual sensors.* The weights of the combination form a spatial filter. Considering *R* ≤ *P* filters, it corresponds to the columns of a matrix **W** ∈ ℝ^*P×R*^. If *R < P*, then spatial filtering reduces the dimension of the data. We will use it here with *R* = *P*. The covariance matrix of ‘spatially filtered’ signals, referred to as source signals, ***W***^⊤^***x**_i_* is readily obtained from ***W**^⊤^**C**_i_**W***. The main idea of the SPoC algorithm is to use the information contained in the outcome variable to guide the decomposition, giving preferences to source signals whose power correlates with *y*. Note that it was originally developed for event-level regression, *e.g.* in BCI, and we adapt it here to a general problem that can also accommodate subject-level regression, where one observation corresponds to one subject instead of one trial. More formally, the filters **W** are chosen to maximize the covariance between the power of the filtered signals and *y*. Denoting by 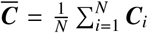 the Euclidean average covariance matrix and 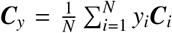 the weighted average covariance matrix, the first filter ***w***_SPoC_ is given by: 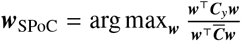. In practice, all the filters in ***W***_SPoC_ are obtained by solving a generalized eigenvalue decomposition problem (Dähne et al., 2014a). Critically we can show that ***W***_SPoC_ recovers the inverse of ***A*** (see proof in Appendix 5.2). Therefore, the SPoC regression model can be defined as a linear model applied to a feature vector of the following form:

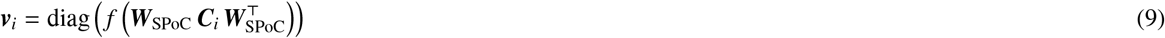

This is consistent for any function *f*. Indeed, since *y_i_* is linearly related to the components of *f* (*p_i_*) that themselves are linearly related to the components of ***v**_i_*, it will also be linearly related to the components of the feature vector ***v**_i_*.

#### Link between the regression models

It is noteworthy that both SPoC and Riemann models have in common to whiten the covariances with a common reference covariance (the Euclidean mean for SPoC and the geometric mean for Riemann): Riemann explicitly with 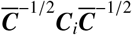, SPoC implicitly by solving generalized eigenvalue problem of 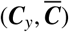, or equivalently of 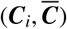 which is equivalent to solving the regular eigenvalue problem of ***C**_i_* after whitening with 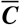 ((Fukunaga, 1990; Nikulin et al., 2011) eq. 13–16). SPoC retrieves the eigenvectors of 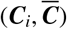. Riemann produces vectors whose size depend on the log eigenvalues of 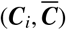. They both produce non-linear features that measure powers relative to a common reference.

‘Upper’ and Riemann models both avoid inverting ***A*** by being insensitive to it. More precisely, they consist in building from ***C_i_*** a *P* × *P* symmetric matrix ***M_i_*** mathematically congruent to a block-diagonal matrix ***D**_i_* whose *Q* × *Q* upper block is diag(*f*(***p***_*i*_)) *i.e.* that writes ***M**_i_* = ***B D***_*i*_ **B**^⊤^, for some invertible matrix ***B***. In this case, the coefficients of *f* (***p_i_***) are a linear combination of coefficients of *M_i_* which implies that the outcome *y_i_* is linear in the coefficients of ***M**_i_*. Therefore a linear model applied to the features ***v**_i_* = Upper(***M**_i_*) is statistically consistent.

Finally, these two models amounts to estimating *Q* parameters (the powers of each sources) from *P*(*P* + 1)/2 parameters (the upper part of a symmetric matrix). Again, it is important to emphasize that we are not aiming for explicitly estimating the most probable model parameters 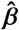 but rather a function that has the smallest approximation error possible, even if over-parametrized. This approach achieves consistency without inverting ***A*** at a price of over-parametrization: the number of parameters will always be a lot higher than the number of samples *N*. Learning in this underdetermined high dimensional setting requires regularizing the linear model to stabilize learning. We will thus use a Ridge regression model with linear kernel, but in a data-driven fashion with nested generalized cross-validation, leading to effective degrees of freedom less than numerical rank of the input data.

In this work we will compare these three models to the inconsistent *diag* model as baseline. This model is probably the historically most frequently used model in M/EEG research in countless publications. Here, powers are considered on the sensor array while the correlation structure is being ignored. This consists in taking only the diagonal elements of the covariance matrix: ***v**_i_* = diag(*f* (***C**_i_*)), which corresponds to the powers (variances) of sensor-level signals.

### 2.5. Model violations

The current theoretical analysis implies that the mixing matrix ***A*** must be common to all subjects and the covariance matrices must be full rank. If these conditions are not satisfied, the consistency guarantees are lost, rendering model performance an empirical question. This will be addressed with simulations (section 2.7), in which we know the true brain-behaviour link, and experiments (section 2.8) in which multiple uncertainties arise at the same time.

#### Individual mixing matrix

A model where the mixing matrix ***A*** is subject-dependent reads: ***x**_i_*(*t*) = ***A**_i_**s***(*t*) + ***n***(*t*). Such situations typically arise when performing subject-level regression due to individual head geometry, individual head positions in MEG and individual locations of EEG electrodes. In this setting, we loose consistency guarantees, but since the ***A**_i_* cannot be completely different from each other (they all originate from human brains), we can still hope that our models perform reasonably well.

#### Rank-deficient signal

In practice M/EEG data is often rank-reduced for mainly two reasons. First, popular techniques for cleaning the data amounts to reduce the noise by projecting the data in a subspace, supposed to predominantly contain the signal. Second, a limited amount of data may lead to poor estimation of covariance matrices: the number of parameters to estimate in a covariance grows quadratically with dimension so many more samples are required than there are sensors to accurately estimate such matrices (Engemann and Gramfort, 2015; Rodrigues et al., 2017). This leads to rank-deficient covariance matrices.

Riemann regression models must be adapted since singular matrices are at infinite distance from any regular matrices. Assuming the rank *R < P* is the same across subjects, the corresponding covariance matrices do not belong to the 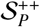 manifold anymore but to the 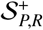 manifold of positive semi-definite matrices of fixed rank *R*. To handle the rank-deficiency one can then project the covariance matrices on 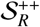 to make them full rank, and then use the geometric distance (See Sabbagh et al. 2019 for more theoretical developments when working with rank-reduced data). To do so, a common strategy is to project the data into a subspace that captures most of the variance. This is achieved by Principal Component Analysis (PCA). We denote the filters in this case by ***W***_UNSUP_ = ***U*** ∈ ℝ^*R×P*^, where ***U*** contains the eigenvectors corresponding to the top *R* eigenvalues of the average covariance matrix 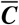. The Riemann regression algorithm is then applied to the spatially-filtered covariance matrix 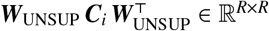.

If covariances are rank-deficient, we will use low-rank versions of both SPoC and Riemann models where only the first components of the spatial filters are kept. In SPoC, components are ordered by covariance with the outcome (supervised algorithm). In Riemann, components are ordered by explained variance in the predictors, not the outcome (unsupervised algorithm). By construction, we can then expect that SPoC achieves similar best performance than Riemann with fewer components: the variance related to the outcome can be represented with fewer dimensions. We can also expect to observe low-rank optima with a plateau after the effective rank *R* of the data.

Besides helping to cope with rank-reduced data, the effect of spatial filtering can be difficult to predict: it helps the regression algorithm by reducing the dimensionality of the problem making it statistically easier, but it can also destroy information if the individual covariance matrices are not aligned (if they span different spaces).

### 2.6. Model-inspection by error-decomposition

The link between the data-generating mechanism and the proposed regression models allows us to derive an informal analysis of variance (Gelman et al., 2005) for estimating the importance of the data generating factors such as head geometry, uniform global power and topographic, i.e., spatial information. Given the known physics from Eq. (1), the data covariance can be written 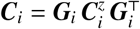, where 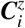 is the covariance matrix of the physiological sources in a given frequency band. The input to the regression model is therefore affected by both the head geometry expressed in ***G**_i_*, and the covariance of the sources. We can therefore compute *degraded* versions of the full covariance 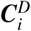 in which only specific components of the signal are retained based on computation of leadfields. Subsequent model comparisons against the full models then allow isolating the relative merit of each component. Following common practice, we considered electrical dipolar sources ***z**_i_*(*t*) ∈ ℝ^*M*^, with *M* ≈ 8000, and we computed the leadfield matrix ***G**_i_* with a boundary element model (BEM) (Gramfort et al., 2014). We then defined two alternative models which are only based on the anatomical information or, additionally, on the global signal power in a given frequency band without topographic structure.

#### Model using anatomy only

Assuming the physiological sources are Gaussian, uncorrelated and of unit variance (power) 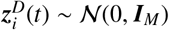, we can re-synthesize their covariance matrix from individual leadfields alone without taking into account the actual covariance structure:

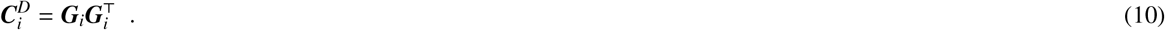

#### Model using anatomy and spatially uniform power

Assuming the physiological sources are Gaussian, uncorrelated and of uniform power 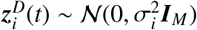 is a scaling factor, we can re-synthesize their covariance matrix from individual leadfields and subject-specific source power, again, ignoring the actual covariance structure:

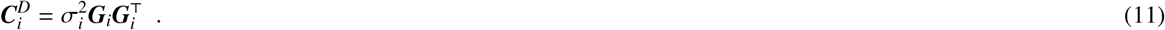

Specifically, we chose 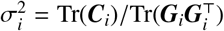, such that 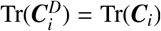: the sum of powers of the signals is the same. This corresponds to taking into account the total power of the sources in a given frequency band and anatomy in the ensuing regression model. Note that we omitted frequency-specific notation for simplicity.

### 2.7. Simulations

We considered simulations to investigate theoretical performance as model violations are gradually introduced. We focused on the ‘linear-in-powers’ and the ‘log-linear-in-powers’ model (Eq. 4 with *f* = identity and *f* = log). Independent identically distributed covariance matrices 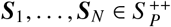 and variables *y*_1_,…, *y_N_* were generated for each generative model. The mixing matrix **A** was defined as exp(*μ**B***) with the random matrix ***B*** ∈ ℝ^*P×P*^ and the scalar *μ* ∈ ℝ to control the distance of *A* from identity (*μ* = 0 yields **A** = ***I**_P_*). The outcome variable was linked to the source powers (i.e. the variance) without and with a log function, and is corrupted by Gaussian noise: *y_i_* = Σ_*j*_ *α_j_f* (*p_ij_* + *ε_i_*, with *f* (*x*) = *x* or log(*x*) and 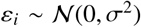 is a small additive random perturbation. We chose *P* = 5, *N* = 100 and *Q* = 2. The affine invariance property of the geometric Riemannian distance should make the model blind to the mixing matrix ***A*** and enable perfect out-of-sample prediction whatever its value when the outcome variable is not corrupted by noise (*σ* = 0). Then we corrupted the clean ground truth data in two ways: by increasing noise in the outcome variable and with individual mixing matrices deviating from a reference: ***A**_i_* = **A** + ***E**_i_*, where entries of ***E**_i_* are i.i.d. 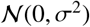. The reference ***A*** can then be thought of as representing the head of a mean-subject.

### 2.8. Experiments: M/EEG data

When analyzing M/EEG data, we do not have access to the actual sources and do not know a priori which generative model, hence, which regression model performs best. Likewise, we cannot expect perfect out-sample prediction: the outcome variable may be noisy (leading to irreducible error), data samples are finite (leading to estimation error) and numerous model violations may apply (mixing matrices may be different for each sample, data may be rank-deficient due to preprocessing, etc.). However, by performing model comparisons based on cross-validation errors, we can potentially infer which model provides the better approximation. We chose three experiments to cover a wide range of model violations. In the order of presentation, they imply rank-deficient covariances (because of limited amount of data) with fixed matrix ***A***, rank-deficient covariances (because of preprocessing) with variable matrices ***A**_i_*, full-rank covariances and variable matrices ***A**_i_*.

#### 2.8.1. Event-level regression: cortico-muscular coherence

We first focused on event-level regression of continuous electromyogram (EMG) from MEG beta activity.

##### Data acquisition

We analyzed one anonymous subject from the data presented in (Schoffelen et al., 2011) and provided by the FieldTrip website to study cortico-muscular coherence (Oostenveld et al., 2011). The MEG recording was acquired with 151 axial gradiometers and the Omega 2000 CTF whole-head system. EMG of forceful contraction of the wrist muscles (bilateral musculus extensor carpi radialis longus) was concomitantly recorded with two silver chloride electrodes. MEG and EMG data was acquired at 1200Hz sampling-rate and online-filtered at 300Hz. For additional details please consider the original study (Schoffelen et al., 2011).

##### Data processing and feature engineering

The analysis closely followed the continuous outcome decoding example from the MNE-Python website (Gramfort et al., 2014). We considered 200 seconds of joint MEG-EMG data. First, we filtered the EMG above 20 Hz using a time-domain firwin filter design, a Hamming window with 0.02 passband ripple, 53 dB stop band attenuation and transition bandwidth of 5Hz (−6dB at 17.5 Hz) with a filter-length of 661 ms. Then we filtered the MEG between 15 and 30 Hz using an identical filter design, however with 3.75 Hz transition bandwidth for the high-pass filter (−6dB at 13.1 Hz) and 7.5 Hz for the low-pass filter (−6dB at 33.75 Hz). The filter-length was about 880 ms. Note that the transition bandwidth and filter-length was adaptively chosen by the default procedure implemented in the filter function of MNE-Python. We did not apply any artifact rejection as the raw data was of high quality. The analysis then ignored the actual trial structure of the experiment and instead considered a sliding window-approach with 1.5 s windows spaced by 250 ms. Allowing for overlap between windows allowed to increase sample size.

We then computed the covariance matrix in each time window and applied Oracle Approximation Shrinkage (OAS) (Chen et al., 2010) to improve conditioning of the covariance estimate. The outcome was defined as the variance of the EMG in each window.

##### Model evaluation

For event-level regression with overlapping windows, we applied 10-fold cross-validation without shuffling such that folds correspond to blocks of neighboring time windows preventing data-leakage between training and testing splits. The initialization of the random number generator used for cross-validation was fixed, ensuring identical train-test splits across models. Note that a Monte Carlo approach with a large number of splits would lead to significant leakage, hence, optimistic bias (Varoquaux et al., 2017). This, unfortunately, limits the resolution of the uncertainty estimates and precludes formalized inference. As we did not have any a priori interest in the units of the outcome, we used the *R*^2^ metric, a.k.a. coefficient of determination, for evaluation.

#### 2.8.2. Subject-level regression with MEG: age prediction

In a second MEG data example, we considered a subject-level regression problem in which we focused on age prediction using the Cam-CAN dataset (Shafto et al., 2014; Taylor et al., 2017).

##### Data acquisition

Cam-CAN dataset contains 643 subjects with resting state MEG data, between 18 and 89 years of age. MEG was acquired using a 306 VectorView system (Elekta Neuromag, Helsinki). This system is equipped with 102 magnetometers and 204 orthogonal planar gradiometers inside a light magnetically shielded room. During acquisition, an online filter was applied between around 0.03 Hz and 1000Hz. To support offline artifact correction, vertical and horizontal electrooculogram (VEOG, HEOG) as well as electrocardiogram (ECG) signal was concomitantly recorded. Four Head-Position Indicator (HPI) coils were used to track head motion. For subsequent source-localization the head shape was digitized. The recording lasted about eight minutes. For additional details on MEG acquisition, please consider the reference publications on the Cam-CAN dataset (Shafto et al., 2014; Taylor et al., 2017).

##### MNE model for regression with source localization

To compare the data-driven statistical models against a biophysics-informed method, for this dataset, we included a regression pipeline based on anatomically constrained minimum norm estimates (MNE) informed by the individual anatomy. Following common practice using the MNE software we used *Q* = 8196 candidate dipoles positioned on the cortical surface, and set the regularization parameter to 1/9 (Gramfort et al., 2014). Concretely, we used the MNE inverse operator as any other spatial filter by multiplying the covariance with it from both sides. We then retained the diagonal elements which provides estimates of the source power. To obtain spatial smoothing and reduce dimensionality, we averaged the MNE solution using a cortical parcellation encompassing 448 regions of interest from Khan et al. (2018). For preprocessing of structural MRI data we used the FreeSurfer software (Fischl 2012, http://surfer.nmr.mgh.harvard.edu/).

##### Data processing and feature engineering

This large dataset required more extensive data processing. We composed the preprocessing pipeline following current good practice recommendations (Gross et al., 2013; Jas et al., 2018; Pernet et al., 2018). The full procedure comprised the following steps: suppression of environmental artifacts, suppression of physiological artifacts (EOG/ECG) and rejection of remaining contaminated data segments.

To mitigate contamination by high-amplitude environmental magnetic fields, we applied the signal space separation method (SSS) (Taulu and Kajola, 2005). SSS decomposes the MEG signal into extracranial and intracranial sources and renders the data rank-deficient. Once applied, magnetometers and gradiometers become linear combinations of approximately 65 common SSS components, hence, become interchangeable (Garcés et al., 2017). For simplicity, we conducted all analyses on magnetometers. We kept the default settings of eight and three components for harmonic decomposition of internal and external sources, respectively, in concert with a 10-second sliding windows (temporal SSS). To discard segments in which inner and outer signal components were poorly separated, we applied a correlation-threshold of 98%. Since no continuous head monitoring data were available at the time of our study, we performed no movement compensation. The origin of internal and external multipolar moment space was estimated based on the head-digitization. To mitigate ocular and cardiac artifacts, we applied the signal space projection method (SSP) (Uusitalo and Ilmoniemi, 1997). This method learns principal components on data-segments contaminated by artifacts and then projects the signal into to the subspace orthogonal to the artifact. To reliably estimate the signal space dominated by the cardiac and ocular artifacts, we excluded data segments dominated by high-amplitude signals using the ‘global’ option from autoreject (Jas et al., 2017). To preserve the signal as much as possible, we only considered the first SSP vector based on the first principal component. As a final preprocessing step, we used the ‘global’ option from autoreject for adaptive computation of rejection thresholds to remove remaining high-amplitudes from the data at the epoching stage.

As the most important source of variance is not a priori known for the problem of age prediction, we considered a wide range of frequencies. We bandpass filtered the data into nine conventional frequency bands *(cf.* Tab. 2) adapted from the Human-Connectome Project (Larson-Prior et al., 2013), and computed the band-limited covariance matrices with the OAS estimator (Chen et al., 2010). All pipelines were separately run across frequencies and features were concatenated after the vectorization step.

**Table 2:** Definition of frequency bands

| name | low | $\delta$ | $\theta$ | $\alpha$ | $\beta_{low}$ | $\beta_{high}$ | $\gamma_{low}$ | $\gamma_{mid}$ | $\gamma_{high}$ |
| --- | --- | --- | --- | --- | --- | --- | --- | --- | --- |
| range (Hz) | 0.1 – 1.5 | 1.5 – 4 | 4 – 8 | 8 – 15 | 15 – 26 | 26 – 35 | 35 – 50 | 50 – 74 | 76 – 120 |

##### Model evaluation

We performed Monte Carlo (shuffle split) cross-validation with 100 splits and 10% testing data. The initialization of the random number generator used for cross-validation was fixed, ensuring identical train-test splits across models. This choice also allowed us to obtain more fine-grained uncertainty estimates than was possible with the time-series data used for subject-level regression. As absolute changes of the unit of the outcome is meaningful, we used the mean absolute error (MAE) as evaluation metric.

#### 2.8.3. Subject-level regression with EEG: age prediction

In a third data example, we considered a subject-level age prediction problem, as in Cam-CAN, but focused on EEG from the Temple University (TUH) EEG Corpus (Harati et al., 2014), one of the largest publicly available database of clinical EEG recordings. This ongoing project currently includes over 30,000 EEGs spanning the years from 2002 to present. This experiment is theoretically important as it supports assessing translation to clinical EEG data. It is also useful as it allows us to evaluate our regression models in a complementary setting where A is sample-dependent, as in Cam-CAN, but covariance matrices are full rank. This experiment is therefore appropriate to primarily investigate the particular model violation of sample-dependent mixing matrices. Unfortunately the absence of associated MRI data prevented us to conduct source localization to correct for individual head geometry.

##### Data acquisition

We used the TUH “Abnormal EEG Corpus”, a subset of TUH EEG Corpus that have been annotated as normal or abnormal by medical experts. From this dataset we focussed on the 1385 healthy patients, from both training and evaluation sets, whose EEG has been annotated as normal. Their age ranges between 10 and 95 years (mean 44.3y, std 16.5y, 775 females, 610 males). EEG was acquired using several generations of Nicolet EEG system (Natus Medical Inc.), equipped between 24 and 36 channels. All sessions have been recorded with an average reference electrode configuration, sampled at 250Hz minimum. The minimal recording length for each session was about 15 minutes. For additional details on EEG acquisition, please consider the reference publications on the TUH dataset (Harati et al., 2014).

##### Data processing, feature engineering and model evaluation

We applied minimal preprocessing to the raw EEG data. We first selected the subset of 21 electrodes common to all subjects (A1, A2, C3, C4, CZ, F3, F4, F7, F8, FP1, FP2, FZ, O1, O2, P3, P4, PZ, T3, T4, T5, T6). We then discarded the first 60 seconds of every recording to avoid artifacts occurring during the setup of the experiment. For each patient we then extracted the first eight minutes of signal from the first session, to be comparable with Cam-CAN. EEG recordings were downsampled to 250Hz. Finally, we excluded data segments dominated by high-amplitude signals using the ‘global’ option from autoreject (Jas et al., 2017) that computes adaptive rejection thresholds. Note that the absence of linear projection to preprocess raw data (as SSS or SSP in Cam-CAN) ensures the data are full rank. While the rank was reduced by one by the use of a common average reference, as we used a subset of channels common to all subjects the data are actually full rank. Otherwise, we followed the same feature engineering and modeling pipeline used for the Cam-CAN data (See section *Subject-level regression with MEG: age prediction*).

#### 2.8.4. Statistical modeling

We used ridge regression (Hoerl and Kennard, 1970) to predict from the vectorized covariance matrices and tuned its regularization parameter by generalized cross-validation (Golub et al., 1979) on a logarithmic grid of 100 values in [10^-5^, 10^3^] on each training fold of a 10-fold cross-validation loop. For each model described in previous sections (‘diag’, ‘upper’, SPoC, Riemann), we standardized the features enforcing zero mean and unit variance. This preprocessing step is standard for penalized linear models. To compare models against chance, we estimated the chance-level empirically through a dummy-regressor predicting the mean of the training data. Uncertainty estimation was obtained from the cross-validation distribution. Note that formal hypotheses testing for model comparison was not available for any of the datasets analyzed as this would have required several datasets, such that each average cross-validation score would have made one observation.

To save computation time, across all analyses, additional shrinkage for spatial filtering with SPoC and unsupervised was fixed at the mean of the value ranges tested in Sabbagh et al. (2019) *i.e.* 0.5 and 10^-5^, respectively.

#### 2.8.5. Software

All analyses were performed in Python 3.7 using the Scikit-Learn software (Pedregosa et al., 2011), the MNE software for processing M/EEG data (Gramfort et al., 2014) and the PyRiemann package (Congedo et al., 2013) for manipulating Riemannian objects. We used the R-programming language and its ecosystem for visualizing the results (Allaire et al., 2019; Clarke and Sherrill-Mix, 2017; R Core Team, 2019; Wickham, 2016). Code used for data analysis can be found on GitHub^1^.

## 3. Results

### 3.1. Simulated data

We simulated data according to the linear and log-linear generative models and compared the performance of the proposed approaches. Fig. 2A displays the results for the linear generative model (*f* = identity in Eq. (4)). The left panel shows the scores of each method as the parameters *μ* controlling the distance from the mixing matrix ***A*** to the identity matrix ***I**_P_* increases (more mixing), and as noise corrupting the outcome variable *y* increases (worse supervision). We see that the Riemannian method is not affected by μ (orange), which is consistent with the affine invariance property of the geometric distance. At the same time, it is not the correct method for this generative model as is revealed by its considerable prediction error greater than 0.5. Unsurprisingly, the ‘diag’ method (green) is highly sensitive to changes in ***A*** with errors proportional to the mixing strength. On the contrary, both ‘upper’ (blue) and SPoC (dark orange) methods lead to perfect out-of-sample prediction (MAE = 0) even as mixing strength increases. This demonstrates consistency of these methods for the linear generative model. They both transparently access the statistical sources, either by being blind to the mixing matrix ***A*** (‘upper’) or by explicitly inverting it (SPoC). Hence, they may enable regression on M/EEG brain signals without source localization. When we add noise in the outcome variable *y* (middle) or individual noise in the mixing matrix (right) we have no theoretical guarantee of optimality for those methods. Yet, we see that both ‘upper’ and SPoC are equally sensitive to these model violations. The Riemannian method seems to be more robust than any other method to individual noise in ***A***, in the sense that its performance is decaying at a slower rate.

**Figure 2:**
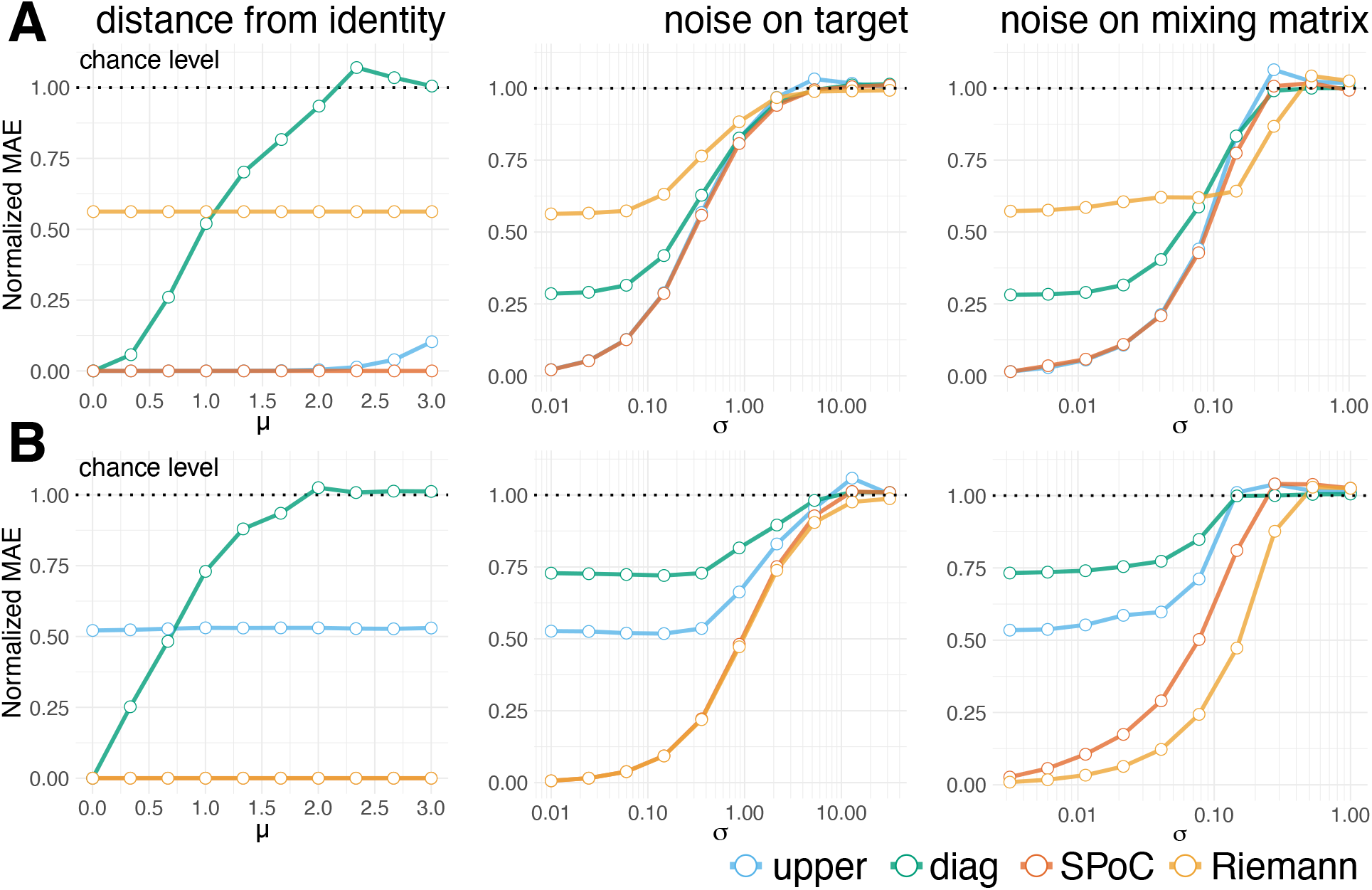
Simulation-based model comparison across generative models. We focused on four regression models (indicated by color) each of which learnt from the covariance in distinct ways. The simulations performance across three types of model violations: the distance ***μ*** between the mixing matrix ***A*** and the identity matrix ***I**_P_* (left), noise on the outcome *y* (middle) and individual noise on ***A*** (right). (**A**) Results for generative model in which y depends linearly on source variance. All but the Riemannian model achieve consistent regression when no mixing occurs (left). SPoC remained consistent throughout the simulated range. The ‘upper’ and SPoC models performed best as noise on the outcome (center) and noise on ***A*** (right) increased. (**B**) Results for generative model in which y depends log-linearly on source powers. The SPoC and Riemannian models achieve consistent regression across all simulated values (left). Likewise, both methods are more robust to noise on *y* (center). Finally, the Riemannian model is most resilient to individual noise on the mixing matrix ***A*** (right).

Fig. 2**B** displays the results for the log-linear generative model (*f* = log in Eq. (4)). In this case Riemann and SPoC performed best (left), as expected by consistency of these methods in this generative model. Both were equally sensitive to noise in outcome variable (middle) but, again, the Riemann method was more robust than other methods as individual noise on the mixing matrix increased (right). The simulations show that, under these idealized circumstances, ‘upper’, and SPoC are equivalent when the outcome y depends linearly on source powers. When y depends linearly on the log-powers, SPoC and Riemann are equivalent. However, when every data point comes with a different mixing matrix, Riemann may be the best default choice, irrespective of the generative model of *y*.

### 3.2. Event-level regression with MEG: Predicting continuous muscle contraction

We then probed the regression models on real MEG data where the true data-generating mechanism is not a priori known and multiple model violations may occur simultaneously. In a first step, we considered a problem where the unit of observation was individual behavior of one single subject with some unknown amount of noise affecting the measurement of the outcome. In this scenario, the mixing matrix is fixed to the extent that the subject avoided head movements, which was enforced by the experimental design. Here, we analyzed one MEG dataset with concomitant EMG recordings that we chose as outcome (See section *Event-level regression: cortico-muscular coherence* in Methods for details). At the time of the analysis, individual anatomical data was not available, hence we constrained the analysis to the sensor-space. The results are depicted in Fig. 3. The analysis revealed that only models including the cross-terms of the covariance predicted visibly better than chance (Fig. 3**A**). For the methods with projection step (SPoC and Riemann) we reported the performance using the full 151 components, equal to the total number of gradiometer channels. Importantly, extensive search for model order for SPoC and Riemann revealed important low-rank optima (Fig. 3**B**) with performance around 50% variance explained on unseen data. This is not surprising when considering the difficulty of accurate covariance estimation from limited data. Indeed, low-rank projection is one important method in regularized estimation of covariance (Engemann and Gramfort, 2015). Interestingly, SPoC showed stronger performance with fewer components than Riemann (4 vs 42). This is not surprising: SPoC is a supervised algorithm, constructed such that its first components concentrate most of the covariance between their power and the outcome variable. The variance related to y can hence be represented with fewer dimensions than Riemann that uses unsupervised spatial filtering. However, it remains equivocal which statistical model best matches this regression problem. The best performing models all implied the log-linear model. Yet, compared to the linear-in-power ‘upper’ model, the low-rank SPoC and Riemann models also implied massive shrinkage on the covariances, leaving unclear if the type of model or regularized covariance estimation explains their superior performance.

**Figure 3:**
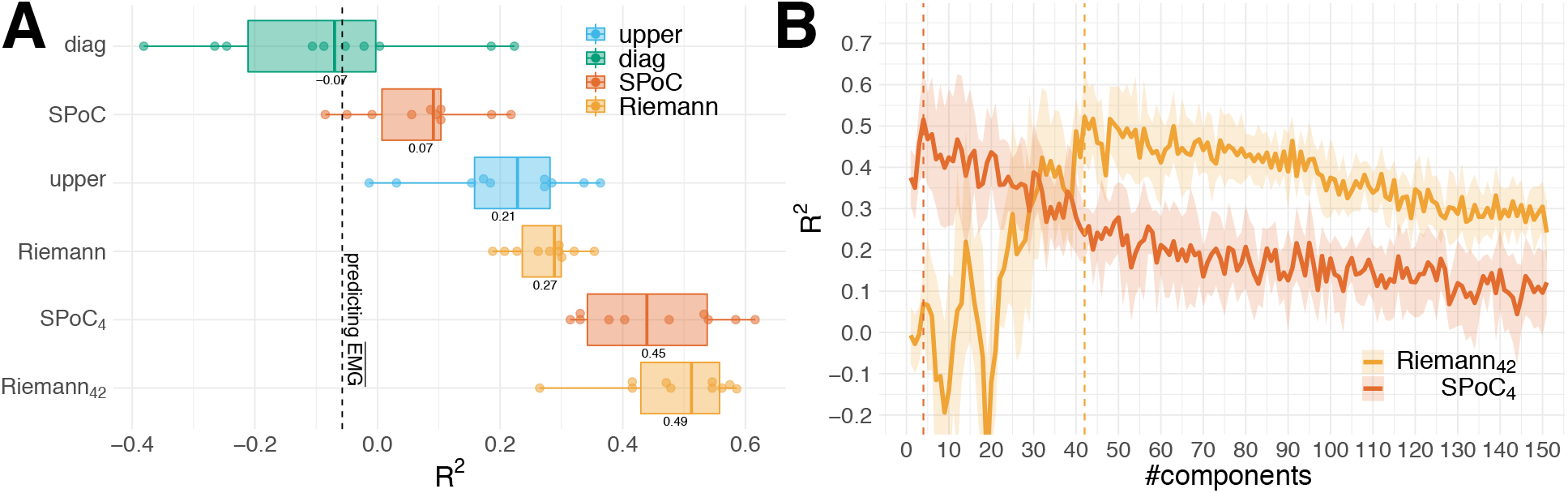
Predicting continuous muscular activity on single-subject MEG. **(A)** model comparison using cross-validation with 10 consecutive groups of approximately overlapping 80 epochs from one single subject. Models are depicted along the y-axis, expected out-of-sample performance (*R*^2^) on the x-axis. The distribution is summarized by standard boxplots. Split-wise prediction scores are represented by dots. The model type is indicated by color. SPoC and Riemann (without subscript) includes spatial filtering with full 151 components, equal to the total number of gradiometer channels. As covariance estimation is necessarily inaccurate with the short 1.5 second epochs, models may perform better when fit on a reduced subspace of the covariance. For these models we reported alternative low-rank models (model order indicated by subscripts). **(B)** Exhaustive search for model order in pipelines with projection step. All values from 1 to the total number of 151 gradiometer channels were considered. One can spot well defined low-rank regimes in both models. However, SPoC supports a lower model order than Riemann. Only models explicitly considering the between-sensor correlation were successful. The best performance was achieved when projecting into a lower dimensional space with optima for SPoC and Riemann of order 4 and 42, respectively.

### 3.3. Subject-level regression with MEG: Predicting age

We then turned our attention to a regression problem that imposes the important model violation of varying source geometry due to individual anatomy while providing a clean outcome with virtually no measurement noise: predicting age from MEG. We analyzed resting-state MEG from about 600 subjects of the Cam-CAN dataset (Shafto et al., 2014; Taylor et al., 2017) focusing on the power spectral topography and between-sensor covariance of nine frequency bands as features (for details see Table 2). In this problem, each sample consists of MEG signals recorded from different persons, hence different brains. On theoretical grounds, one may therefore expect individual cortical folding, size and proportional composition of the head and its tissues to induce important distortions to the signal that may pose severe problems to purely data-driven approaches. Here, each data point can be said to have its own mixing matrix inducing unique distortions in each observation. To investigate this point explicitly, we conducted source localization to obtain power estimates that corrected for individual head geometry based on biophysical prior knowledge. On the other hand, 8 minutes of MEG support accurate covariance estimation, hence, rendering model order search less important for shrinkage. Covariance matrices are nevertheless rank-deficient due to SSS and SSP preprocessing steps. Fig. 4 displays the results for different regression models.

**Figure 4:**
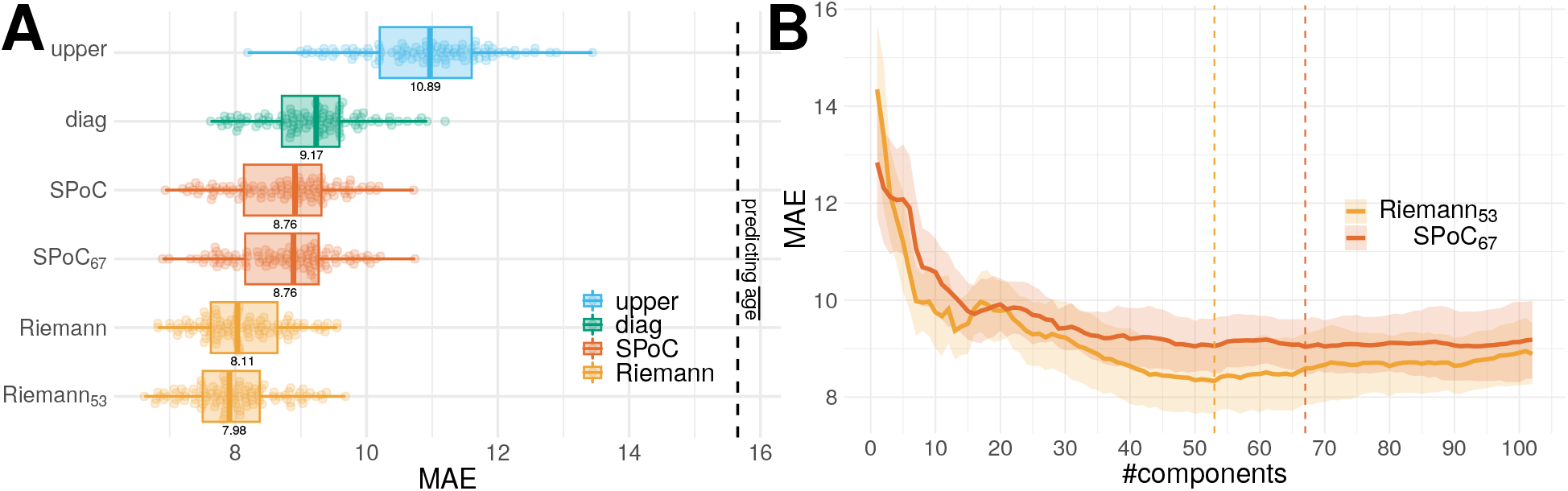
Predicting age from subject-level MEG in sensor space. **(A)** model comparison using Monte Carlo cross-validation with 100 splits sampled from 596 subjects. Models are depicted along the y-axis, expected out-of-sample performance (mean absolute error) on the x-axis. The distribution is summarized by standard boxplots. Split-wise prediction scores are depicted by dots. The model type is indicated by color. Here, covariance estimation was based on 8 minutes of MEG, hence, the impact of shrinkage should be small. For comparison with the single-subject data (Fig. 3), we nevertheless reported the alternative low-rank models (model order indicated by subscripts, no subscript meaning an order of 65, the minimum rank of covariances). **(B)** Exhaustive search for model order in pipelines with projection step. All values from 1 to the total number of 102 magnetometer channels were considered. One can see that performance starts to saturate around 40 to 50. No striking advantage of model order search was evident compared to deriving the order from prior knowledge on rank deficiency at a value of about 65. All models performed better than chance, however, models consistent with log-linear model and using correlation terms performed better. The Riemannian models performed best.

The analysis revealed that all models performed clearly better than chance. The Riemannian model (orange) yielded the best performance (8y MAE), followed by SPoC (dark orange, 8.8y MAE) (4**A**). The diagonal (green) and upper-triangle (blue) models performed worse. Model order search did not reveal striking low-rank optima. Models above rank 40 seem approximately equivalent, especially when considering the estimation uncertainty of standard deviation above 1 year of MAE. For both SPoC and Riemann, the best low-rank model was close to the model at the theoretically derived rank of 65 (due to preprocessing with SSS, see Taulu and Kajola 2005). For subsequent analyses, we, nevertheless, retained the best models.

One first important observation suggests that the log-linear model is more appropriate in this regression problem, as the only model not implying a log transform, the ‘upper’ model, performed clearly worse than any other model. Yet, important difference in performance remain to be explained among the log-linear models.

This points at the cross-terms of the covariance, which turns out to be an essential factor for prediction success: The ‘diag’ model ignores the cross-terms and performed worst among all log-linear models. The SPoC and Riemann models performed better than ‘diag’ and both analyzed the cross-terms, SPoC implicitly through the spatial filters. This raises the question why the cross-terms were so important. One explanation would be that they reveal physiological information regarding the outcome. Alternatively, the cross-terms may expose the variability due to individual head geometry. To further investigate this point we conducted the same regression analysis on source localized M/EEG signals, i.e., after having corrected for individual head geometry with a biophysical model.

#### Source space analysis

Next, we have computed source-space covariance matrices based on anatomically constrained Minimum Norm Estimates (MNE). Results are depicted in Fig. 5. Now, the optimal number of components for prediction remarkably dropped: 11 for Riemann and 20 for SPoC in source space, as compared to 53 and 67, respectively, in sensor space. This may suggest that the inflated number of components in sensor space is related to extra directions in variance accounting for individual head geometry. Second, ‘diag’ (green) is now the best regression model. This model only takes the log powers into account and discards the cross-terms. This suggests that the outcome does not depend on the cross-terms or at least that the potential gain of the cross-terms is inaccessible due to the inflated dimensionality of feature space. The ‘diag’ score is also the highest among all the models that we considered so far, illustrating that the MNE solution to the inverse problem provides superior unmixing of brain signals.

**Figure 5:**
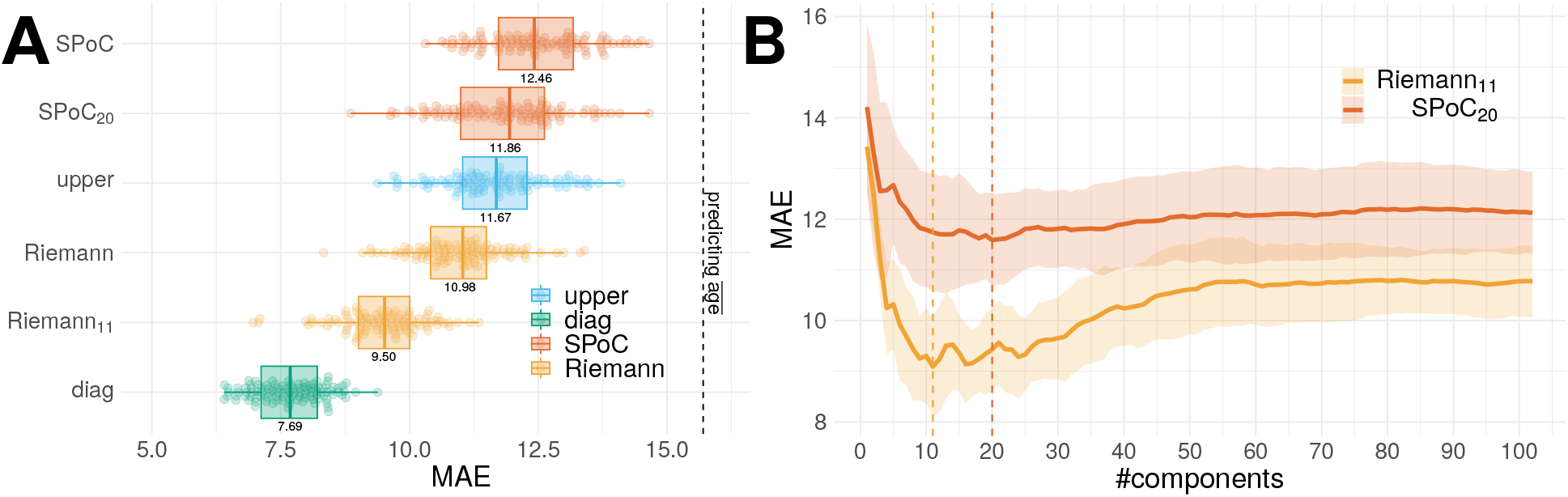
Predicting age from subject-level MEG in source space. **(A)** model comparison applied to sources and using Monte Carlo crossvalidation with 100 splits sampled from 596 subjects. It follows the same layout conventions than Fig. 4. The sources are estimated by MNE that exploits biophysical prior knowledge. **(B)** Exhaustive search for model order in pipelines with projection step. All values from 1 to the total number of 102 magnetometer channels were considered. One can see that performance starts to saturate around 40 to 50. But contrary to sensor space analysis of Fig. 4 projection models show clear low-rank minima. All models performed better than chance, however, the ‘diag’ model that only considers sources’ log-powers clearly outperforms other models.

The performance of Riemann in sensor space is, nevertheless, close to ‘diag’ in source space, suggesting that the cross-term models, in sensor space, have learnt to some extent what ‘diag’, in source space, receives explicitly from source localization. Still, the good performance of ‘diag’ in source space may be due to two independent factors that are not mutually exclusive: It could be that source localization standardizes head geometry, hence, mitigates the variability of mixing. On the other hand, if the anatomy itself covaries with the outcome, which is a safe assumption to make for the case of aging (Liem et al., 2017), the leadfields will also covary with the outcome. Source amplitudes may then change as a result of dampening-effects (See methods in Khan et al. (2018)). To investigate these factors, we conducted an error-decomposition experiment.

#### Error decomposition

To disentangle the factors explaining model performance, we devised a novel error-decomposition method derived from the proposed statistical framework (Fig. 1). Using a simulation-based approach, we computed degraded observations, *i.e.*, individual covariance matrices, that were either exclusively influenced by the individual anatomy in terms of the leadfields (Eq. 10) or also by additive uniform power (Eq. 11). For details see section *Model-inspection by error-decomposition* in Methods. This has allowed us to estimate to which extent the log-linear models have learnt from anatomical information, global signal power of the MEG and topographic details.

Fig. 6 compares three log-linear models based on the original observations (black) and the degraded covariances (orange): the ‘diag’ model and the best low-rank models previously found for SPoC and Riemann methods.

**Figure 6:**
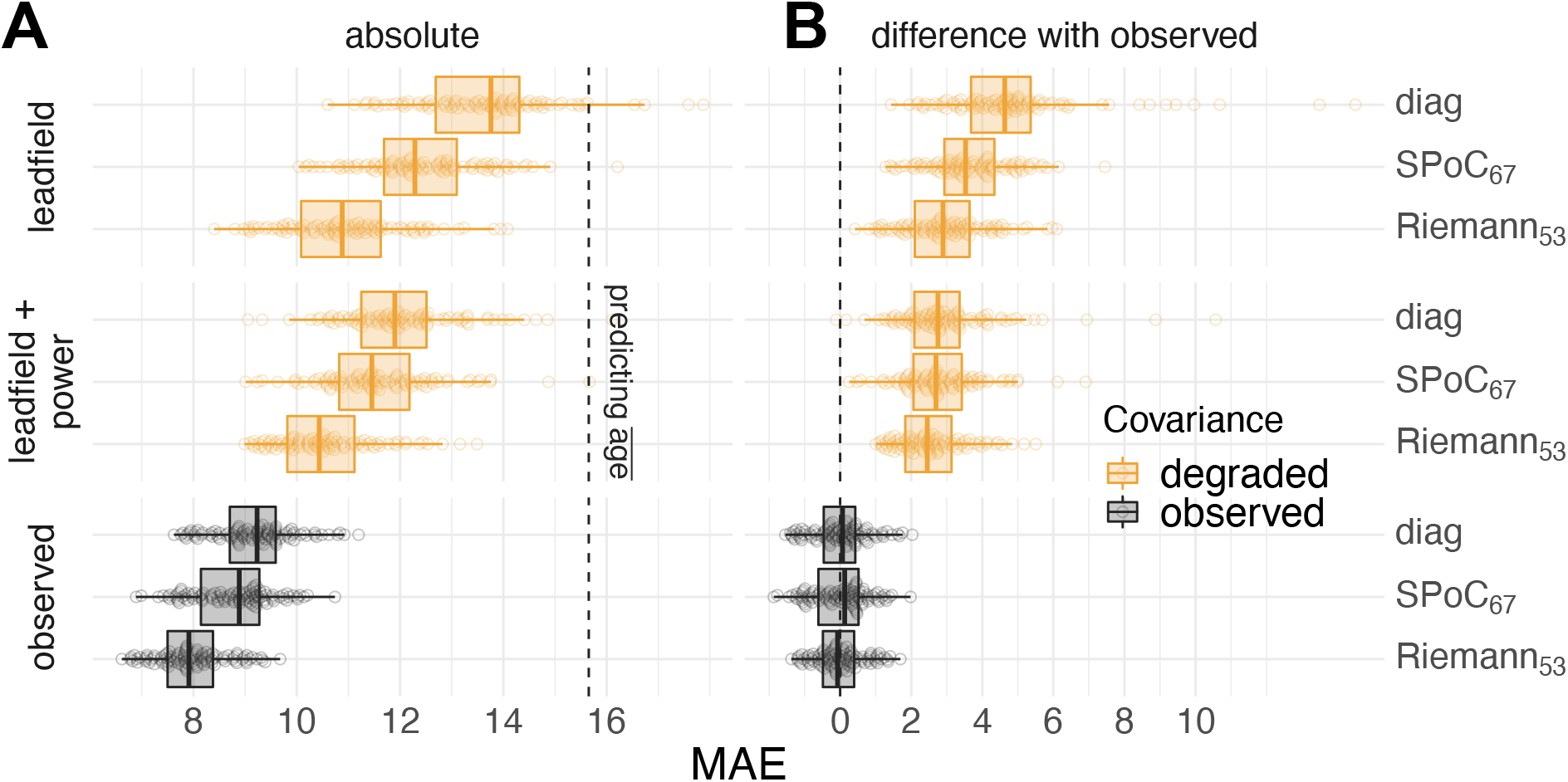
Simulation-based error decomposition. We performed model comparisons for the observed data (black) and degraded data (orange) for which spatio-spectral information was progressively removed: ‘leadfield + power’ muted topographic information keeping only spatially uniform power and information from the individual leadfields (Eq. 11), ‘leadfield’ muted all electrophysiological variance (Eq. 10). (**A**) depicts absolute performance, (**B**), differences with the full observation, correspondingly, for each model. One can see that all models learnt to predict age from all three components: anatomical variation across subjects, electrophysiological signal power and topographic information. However, the relative importance of each error component was clearly different across models. Riemannian model was most responsive to the leadfield component and least responsive to the uniform power.

One can see that all three error components improved overall prediction in similar ways, each improving performance between 2 and 4 years on average (Fig. 6**A**). The best performance with the leadfields-only was obtained by the Riemannian model scoring an MAE of about 11 *y* on average. Adding spatially uniform power, the Riemann model kept leading and improved by about 0.5*y*. Predictions based on the observed data with full access to the covariance structure improved performance by up to about 3*y*, suggesting that age prediction clearly benefits from information beyond the leadfields.

Generally, the choice of algorithm mattered across all levels of the data generating scenario with Riemann always leading and the ‘diag’ model always trailing (Fig. 6**A**). Finally, the results suggest the presence of an interaction effect where both the leadfields and the uniform power components were not equally important across models (Fig. 6**A,B**). For the Riemannian model, when only learning from leadfields, performance got as close as three years to the final performance of the full model (Fig. 6**B**). The ‘diag’ model, instead, only arrived at 5 years of distance from the equivalent model with full observations (Fig. 6**B**). On the other hand, the Riemannian model extracted rather little additional information from the uniform power and only made its next leap forward when accessing the full nondegraded covariance structure. Please note that these analyses are based on cross-validation. The resulting resampling splits do not count as independent samples. This precludes formal analysis of variance with an ANOVA model.

Overall, error decomposition suggests that all methods learn from anatomy and that indeed, the leadfield in isolation is predictive of age. Models considering cross-terms of the covariance were however more sensitive.

#### Robustness to preprocessing choices

This leads to a final consideration about error components. Commonly used preprocessing in M/EEG analysis is based on the idea to enhance signal-to-noise ratio by removing signals of noninterest, often using dedicated signal-space decomposition techniques (Hyvärinen et al., 2004; Taulu and Kajola, 2005; Uusitalo and Ilmoniemi, 1997). However, it is perfectly imaginable that such preprocessing removes information useful for predicting. At the same time, predictive models may learn the signal subspace implicitly, which could render preprocessing unnecessary. To investigate this issue for the current subject-level regression problem, we sequentially repeated the analysis after activating the essential preprocessing steps one by one, and compared them to the baseline of extracting the features from the raw data. For this purpose, we considered an alternative preprocessing pipeline in which we kept all steps unchanged but the SSS (Taulu and Kajola, 2005) for removal of environmental artifacts. We used instead a data-driven PCA-based SSP (Uusitalo and Ilmoniemi, 1997) computed on empty room recordings. The preprocessing pipeline is detailed in section *Subject-level regression with MEG: age prediction.* Results are depicted in Fig. 7.

**Figure 7:**
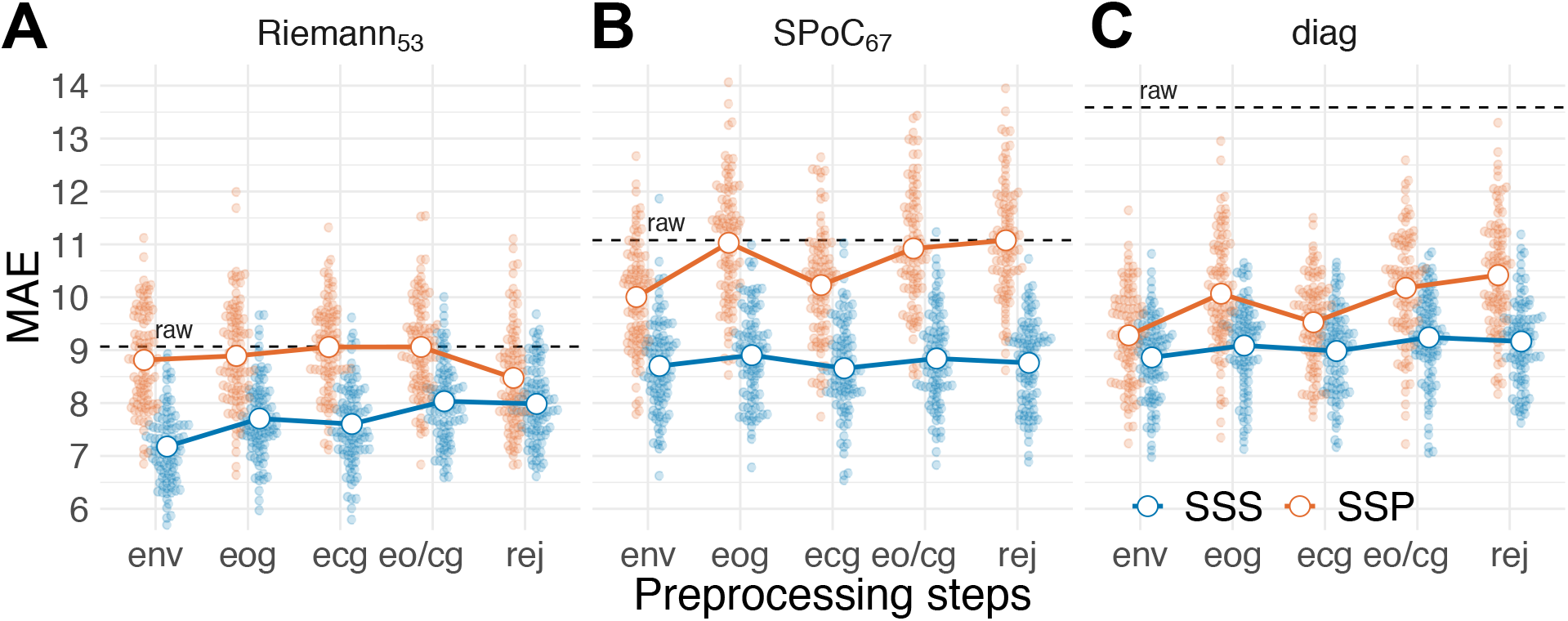
Impact of preprocessing. Model comparison across cumulative artifact removal steps: environmental artifacts (env), environmental + occular (eog), environmental + cardiac (ecg), environmental + occular + cardiac (eo/cg), environmental + occular + cardiac + bad segments (rej). Results are compared to the baseline of extracting features from raw data with no preprocessing (depicted by vertical dashed lines). The method for removal of environmental artifacts is indicated by color, *i.e.*, blue and red for SSS and SSP respectively. Note that the endpoint rej is identical to the full preprocessing conducted in previous analyses. Panels depict performance for the best Riemannian model **(A)**, the best SPoC model **(B)**, and the ‘diag’ model **(C)**. One can see that the Riemann model, but not the ‘diag’ model, is relatively robust to preprocessing and its details.

The analysis revealed that the Riemannian model performed reasonably well when no preprocessing was done at all (Fig. 7**A**). It also turned out to be relatively robust to particular preprocessing choices. On the other hand, whether preprocessing was done or not turned out decisive for the ‘diag’ model and to some extent for the SPoC model (Fig. 7**B,C**). A few common tendencies became apparent. Across all models, while improving above baseline, SSP consistently led to worse performance than SSS. Second, performance was also slightly degraded by removing ocular and cardiac artifacts, suggesting that both shared variance with age. Removing EOG seemed to consistently degrade performance. On the other hand, removing ECG had virtually no impact for SPoC and the ‘diag’ model. For Riemann, both removing ECG and EOG additively deteriorated performance. Finally bad epochs rejection had a negligible and inconsistent effect. Overall, the results suggest that the importance of preprocessing depended on the model, while minimal denoising with SSP or SSS always helped improve performance. Of note, with minimal preprocessing using SSS, the Riemannian model performed at least as well as the ‘diag’ model after source localization (Fig. 5).

### 3.4. Subject-level regression with clinical EEG: Predicting age

The results on subject-level regression based on MEG suggest the importance of model violations due to individual head geometry. Importantly, with traditional cryogenic MEG, the sensor array is not fixed relative to the head, rendering head-positioning and head-movements factors contributing to model violations due to individual signal geometry. How would the present results generalize to clinical EEG where sensors are fixed relative to the head but, in general, fewer sensors are used ? To investigate this question, we applied our subject-level regression setting to clinical EEG: We analyzed resting-state EEG (21 sensors) from about 1000 subjects of the TUH dataset (Harati et al., 2014). As with previous analysis of the Cam-CAN data, each data point had its own mixing matrix. As with the Cam-CAN, the EEG recordings from TUH were sufficiently long to support accurate covariance estimation, hence, rendering model order search less important for shrinkage. We did not preprocess the data on purpose to ensure having full-rank signals. This experiment is therefore appropriate to primarily investigate the particular model violation of sample-dependent mixing matrices with constrained degrees of freedom for the sensor-positioning as well as the generalization from MEG to EEG. Fig. 8 displays the results for different regression models. Unfortunately the absence of associated MRI data prevented us to conduct source localization to correct for individual head geometry.

**Figure 8:**
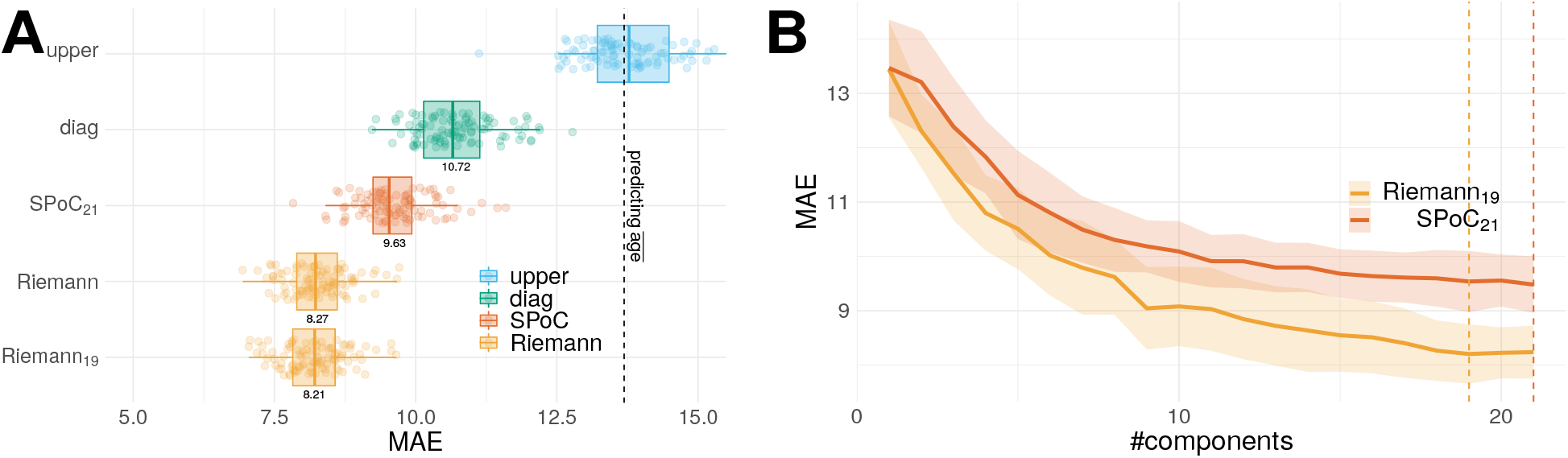
Predicting age from subject-level EEG in sensor space. **(A)** model comparison applied to sensors and using Monte Carlo crossvalidation with 100 splits sampled from 1000 subjects. It follows the same layout conventions than Fig. 4. Here, covariance are full rank, the impact of shrinkage should be small. We nevertheless reported the alternative low-rank models (model order indicated by subscripts). **(B)** Exhaustive search for model order in pipelines with projection step. All values from 1 to the total number of 21 electrodes were considered. Model order search did not reveal striking low-rank optima. All models except ‘upper’ performed better than chance, however, models consistent with log-linear model and using correlation terms performed better. The Riemannian models performed best.

Model order search did not reveal any clear low-rank optima. This was expected considering the absence of preprocessing and accurate covariance estimation. Strikingly, the only model not implementing a log transform, the ‘upper’ model, performed at chance level, clearly worse than any other model. All other models performed better than chance, with Riemann clearly leading, followed by SPoC and diag. Those results are consistent with our simulations in Fig. 2**(B)** in which the only model violation comes from individual mixing matrices. The performance and ordering of the models in the TUH data is also consistent with the results obtained on the Cam-CAN dataset. This strongly suggests that the log-linear model is more appropriate in this regression problem. It is noteworthy, that the best performance based on the Riemannian model was virtually identical to its performance with MEG on the Cam-CAN data. However, it remains open to which extent the benefit of constrained signal geometry due to fixed sensor positioning is cancelled out by reduced spatial sampling with 21 instead of 306 sensors.

## 4. Discussion

In this work, we have proposed a biophysics-aware framework for predictive modeling with M/EEG. We specifically considered regression tasks based on source power in which source localization is not practical. To the best of our knowledge, this is the first study systematically comparing alternative approaches for predicting continuous outcomes from M/EEG in a coherent theoretical framework from the level of mathematical analysis over simulations down to analysis of real data. Here, we focused on the band-limited between-channels covariance as fundamental representation and investigated distinct regression models. Mathematical analysis identified different models supporting perfect prediction under ideal circumstances when the outcome is either linear or log-linear in the source power. We adapted techniques originating from event-level prediction *i.e.* the SPoC spatial filtering approach (Dähne et al., 2014a; de Cheveigné and Parra, 2014) and projection with Riemannian geometry (Congedo et al., 2017) for subject-level prediction typically encountered in biomarker development. Our simulation-based findings were consistent with the mathematical analysis and suggested that the regression models based on adaptive spatial filtering or Riemannian geometry were more robust across data generating scenarios and model violations. Subsequent analysis of MEG focused on a) differences between models in event-level and subject-level prediction, b) the contribution of anatomical and electrophysiological factors, for which we proposed a novel error decomposition method, and c) the impact of preprocessing, investigated through large-scale analysis of M/EEG data. Our findings suggest that, consistent with simulations, Riemannian methods are generally a good bet across a wide range of settings with considerable robustness to different choices of preprocessing.

### 4.1. What distinguishes event-level from subject-level prediction in the light of model violations?

Unsurprisingly, no model performed perfectly when applied to empirical data for which the data generating mechanism is by definition unobservable, multiple model violations may occur and information is only partially available. One important source of differences in model violation is related to whether outcomes are defined at the event-level or at the subject-level. When predicting outcomes from ongoing segments of neural time-series within a subject, covariance estimation becomes non-trivial as the event-level time windows are too short for accurate estimation. Even if regularized covariance estimates provide an effective remedy, there is not one shrinkage recipe that works in every situation (Engemann and Gramfort, 2015). In this study, we have relied on the oracle approximating shrinkage (OAS) (Chen et al., 2010) as a default method in all analyses. Yet, we found that additional low-rank shrinkage (Engemann and Gramfort, 2015; Rodrigues et al., 2018; Tipping and Bishop, 1999; Woolrich et al., 2011), as implied by the SPoC method (Dähne et al., 2014a), or the unsupervised projection for the Riemannian model (Sabbagh et al., 2019), improved performance considerably for event-level prediction. A spatial-filter method like SPoC (Dähne et al., 2014a; de Cheveigné and Parra, 2014) can be particularly convenient in this context. By design, it concentrates the variance most important for prediction on a few dimensions, which can be easily searched for, ascending from the bottom of the rank spectrum. Riemannian methods can also be operated in low-rank settings (Sabbagh et al., 2019). However, model order search may be more complicated as the best model may be anywhere in the spectrum. This can lead to increased computation times, which may be prohibitive in realtime settings such as BCI (Lotte et al., 2018, 2007; Tangermann et al., 2008).

Issues with the numerical rank of the covariance matrix also appear when predicting at the subject-level. The reason for this is fundamentally different and rather unrelated to the quality of covariance estimation. Many modern M/EEG preprocessing techniques focus on estimating and projecting out the noise-subspace, which leads to rank-deficient data. In our analysis of the Cam-CAN dataset (Shafto et al., 2014; Taylor et al., 2017), we applied the SSS method (Taulu and Kajola, 2005) by default, which is the recommended way when no strong magnetic shielding is available, as is the case for the Cambridge MEG-system on which the data was acquired (see also discussion in Jas et al. 2018). However, SSS massively reduces the rank down to about 64 out of 306 dimensions, which may demand special attention when calibrating covariance estimation. Our results suggest that projection can indeed lead to slightly improved average prediction once a certain rank value is reached. Yet, thoughtful search of optimal model order may not be worth the effort in practice when a reasonably good guess of model order can be derived from the understanding of the preprocessing steps applied. Our findings, moreover, suggest, that a Riemann-based model is, in general, a reasonably good starting point, even when no model order search is applied. What seems to be a much more important issue in subject-level prediction from M/EEG are the model violations incurred by individual anatomy. Our mathematical analysis and simulations demonstrated that not even the Riemannian approach is immune to those, for MEG and EEG.

### 4.2. What explains the performance in subject-level prediction?

Our results suggested that, for the current regression problems with MEG and EEG, the log-linear model was more appropriate than the linear-in-powers ones. This is well in line with practical experience and theoretical results highlighting the importance of log-normal brain dynamics (Buzsáki and Mizuseki, 2014). On the other hand, on the Cam-CAN data, we observed substantive differences in performance within the log-normal models highlighting a non-trivial link between the cross-terms of the covariance and subject-level variation. Indeed, the ‘diag’ model, both in sensor and source space, ignored the cross-terms of the covariance, yet in source space, it performed about 1.5 years better on average than in sensor space. This is rather unsurprising when recapitulating the fact that subject-level regression on M/EEG implies individual anatomy. Indeed, our mathematical analysis and simulations identified this factor as important model violation. MNE source localization, by design, uses the head and brain geometry to correct for such violations. On the other hand, if leadfields are correlated with the outcome, the source localization, which depends on the leadfields, will be predictive of the outcome too, even if no brain source is actually relevant to the outcome. This suggests that the cross-term models that were more successful than the ‘diag’ model may either convey biological information relevant to predict the outcome, or expose forward information on head geometry to the regression model, which then improved prediction by de-confounding for head geometry. Our findings on source localization strongly suggested that correcting for geometrical misalignment was the driving factor, evidenced by the fact that after source localization the simple ‘diag’ model performed best. Yet, these findings did not rule out that leadfields themselves were not predictive of the outcome.

We, therefore, derived a novel error-decomposition technique from the statistical framework presented in Fig. 1 to estimate the sensitivity of our M/EEG regression models to anatomy, spatially uniform power and topographic details. We applied this technique on the Cam-CAN dataset to investigate the subject-level prediction problem. While all models captured anatomical information and the Riemannian models were the most sensitive to it, anatomical information did not explain the performance based on the full data. At the same time, this demonstrated that MEG captures age-related anatomical information from the individual leadfields and raises the question of which aspects of anatomy were concerned. Neuroscience of aging has suggested important alterations of the cortical tissues (Liem et al., 2017), relevant for generating M/EEG signals, such as cortical surface area, cortical thickness or cortical folding. Yet, more trivially, head size or posture are a common issue in MEG and could explain the present effect, which would be potentially less fascinating from a neuroscientific standpoint. We investigated this issue post-hoc by predicting age from the device-to-head transform describing the position of the head relative to the helmet and the coregistration transforms from head to MRI. Compared to the Riemannian model applied to the leadfields-only surrogate data, this resulted in three years lower performance of around 14 years error, which is close to the random guessing error and may at best explain the performance of the ‘diag’ model. Moreover, translating our approach to EEG for which sensor placement relative to the head is less variable, we did not witness improvements over MEG. On the other hand, this may be due to the smaller number of sensors available in EEG. Future work will have to show, how these two factors interact in practice across prediction problems and EEG-configurations.

Interestingly, also the SPoC model was more sensitive to anatomy than the ‘diag’ model. This suggests that by learning adaptive spatial filters from the data to best predict age, SPoC may implicitly also tune the model to the anatomical information conveyed by the leadfields. This seems even more plausible when considering that from a statistical standpoint, SPoC learns how to invert the mixing matrix ***A*** to get the statistical sources implied by the predictive model. This must necessarily yield a linear combination of the columns of ***G***. As a consequence, SPoC does not learn to invert the leadfields ***G*** but directly yields an imperfect approximation to ***G***. Theoretically, unique SPoC solution can be found with arbitrary outcomes as long as the data is full-rank and the target is noise-free. In practice, this is rarely the case. Therefore, the SPoC solution empirically depends on the choice of the outcome. This also motivates the conjecture that differences between SPoC and Riemann should become smaller when the ***G**_i_*, are not correlated with the outcome (Riemann should still enjoy an advantage due to increased robustness to model violations) or even vanish when ***G*** is constant and no low-rank issues apply. The latter case is what we encountered in the event-level analysis where SPoC and Riemann where roughly on par, suggesting that both handled the distortions induced by ***G***.

Unfortunately, the current analysis did not elucidate the precise mechanism by which different models learnt from the individual anatomy and why the Riemannian model was so much more proficient. As a speculation, one can imagine that changes in the leadfields translate into simple topographic displacements that the ‘diag’ model can easily capture. This would be in line with the performance of the ‘diag’ model on the leadfields-only surrogate data, which matched prediction performance based on the device-to-head transforms or the coregistration matrices previously mentioned. With cross-terms included in the modeling, SPoC and, in particular, Riemann may better unravel the directions of variation with regard to the outcome by considering the entire geometry presented in the leadfields. Instead, for the case of the leadfields-only surrogates, SPoC attempts capturing sources which literally do not exist, hence must yield a degraded view on ***G***.

Overall, our results suggest that Riemannian models may also be the right choice when the anatomy is correlated with the outcome and the primary goal is prediction. The enhanced sensitivity of the Riemannian model to source and head geometry may be precisely what brings them so close to performance based on source localization. Indeed, the TUH experiment shows that these properties render Riemannian models particularly helpful in the case of EEG, where the leadfields should be less variable as the sensor cap is affixed to the head, which strongly limits variation due to head posture.

### 4.3. How important is preprocessing for subject-level prediction?

It is up to now equivocal how important preprocessing is when performing predictive modeling at the subject-level. Some evidence suggests that preprocessing may be negligible when performing event-level decoding of evoked responses as a linear model may well learn to regress out the noise-subspace (Haufe et al., 2014b). Our findings suggest a more complex situation when performing subject-level regression from M/EEG signal power. Strikingly, performing no preprocessing was clearly reducing performance, for some models even dramatically, SPoC and in particular ‘diag’. The Riemann model, on the other hand, was remarkably robust and performed even reasonably well without preprocessing. Among the preprocessing steps, the removal of environmental artifacts seemed to be most important and most of the time led to massive improvements in performance. Removing EOG and ECG artifacts mostly reduced performance suggesting that age-related information was present in EOG and ECG. For example, one can easily imagine that older subjects produced less blinks or showed different eye-movement patterns (Thavikulwat et al., 2015) and also cardiac activity may change across the lifespan (Attia et al., 2019).

Interestingly, our results suggest that the method used for preprocessing was highly important. In general, performance was clearly enhanced when SSS was used instead of SSP. Does this mean that SSP is a bad choice for removing environmental artifacts? Our results have to be interpreted carefully, as the situation is more complicated when considering how fundamentally different SSP and SSS are in terms of design. When performing SSS, one actually combines the information of independent gradiometer and magnetometer sensor arrays into one latent space of roughly 65 dimensions, less than the dimensionality of both sensor arrays (306 sensors in total). Even when analyzing the magnetometers only after SSS, one will also access the extra information from the gradiometers (Garcés et al., 2017). SSP on the other hand is less invasive and is applied separately to magnetometers and gradiometers. It commonly removes only few dimensions from the data, yielding a subspace greater than 280 in practice. Our results therefore conflate two effects: 1) learning from magnetometers and gradiometers versus learning from magnetometers only and 2) differences in strength of dimensionality reduction. To disentangle these factors, careful experimentation with more targeted comparisons is indicated. To be conclusive, such an effort may necessitate computations at the scale of weeks and should be investigated in a dedicated study. For what concerns the current results, the findings simply suggest that SSS is a convenient tool as it allows one to combine information from magnetometers and gradiometers into a subspace that is sufficiently compact to enable efficient parameter estimation. It is not clear though, if careful processing with SSP and learning on both sensors types would not lead to better results.

## 5. Conclusion

Our study has investigated learning continuous outcomes from M/EEG signal power from the perspective of generative models. Across datasets and electrophysiological modalities, the log-linear model turned out to be more appropriate. In the light of common empirical model violations and preprocessing options, models based on Riemannian geometry stood out in terms of performance and robustness. The overall performance level is remarkable when considering the simplicity of the model. Our results demonstrate that a Riemannian model can actually be used to perform end-to-end learning (Schirrmeister et al., 2017) involving nothing but signal filtering and covariance estimation and, importantly, without deep-learning (Roy et al., 2019). When using SSS, performance improves beyond the current benchmark set by the MNE model but probably not because of denoising but rather due to the addition of gradiometer information. Moreover, we observed comparable performance on minimally processed clinical-EEG with only 21 channels instead of 306 MEG-channels, suggesting that the current approach may well generalize to certain clinical settings. This has at least two important practical implications. First, this allows researchers and clinicians to quickly assess the limits of what they can hope to learn in an economical and eco-friendly fashion (Strubell et al., 2019). In this scenario, the Riemannian end-to-end model rapidly delivers an estimate of the overall performance that could be reached by extensive and long processing, hence, support practical decision making on whether a deeper analysis is worth the investment of time and resources. Second, this result suggests that if prediction is the priority, availability of MRI and precious MEG expertise for conducting source localization is not any longer the bottleneck. This could potentially facilitate data collection and shift the strategy towards betting on the law of large numbers: assembling an MEG dataset in the order of thousands is easier when collecting MRI is not a prerequisite.

It is worthwhile to consider important limitations of this study. Unfortunately, we have not had access to more datasets with other interesting continuous outcomes. In particular the conclusions drawn from the comparison between event-level and subject-level regression may be expanded in the future when considering larger event-level datasets and other outcomes for which the linear-in-powers model may be more appropriate. Second, one has to critically acknowledge that the performance benefit for the Riemannian model may be partially explained by increased sensitivity to anatomical information, which might imply reduced specificity with regard to neuronal activity. In this context it is noteworthy that recent regression pipelines based on a variant of SPoC (Dähne et al., 2014b) made use of additional spatial filtering for dimensionality reduction, *i.e.*, SSD (Nikulin et al., 2011) to isolate oscillatory components and discard arrhythmic (1/f) activity. This raises the question if the specificity of a Riemannian model could be enhanced in a similar way. Ultimately, what model to prefer, therefore, clearly depends on the strategic goal of the analysis (Bzdok et al., 2018; Bzdok and Ioannidis, 2019) and cannot be globally decided.

We hope that this study will provide the community with the theoretical framework and tools needed to deepen the study of regression on neural power spectra and safely navigate between regression models and geometric distortions governing M/EEG observations.

## Acknowledgements

This work was supported by a 2018 “médecine numérique” (for digital medicine) thesis grant issued by Inserm (French national institute of health and medical research) and Inria (French national research institute for the digital sciences). It was also partly supported by the European Research Council Starting Grant SLAB ERC-StG-676943. The authors would like to thank Lucas C. Parra and Marijn van Vliet for critical reviewing and suggestions on earlier versions of the manuscript. The authors would also like to thank Lukas Gemein and Tonio Ball from the Neuromedical AI Lab at University of Freiburg for their help in extracting the age from the TUH dataset. The the authors thank Sheraz Khan for help with the Freesurfer segmentation and data management of the Cam-CAN dataset. The the authors thank Donald Krieger and Timothy Bardouille for help with the MEG co-registration.

## Appendix

### 5.1 Full-rank formulation of M/EEG signal generative model

Denoting by 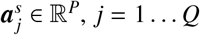 the independent *source patterns*, the generative model of M/EEG observations reads:

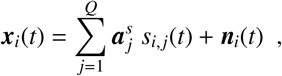

where ***s**_i,j_*(*t*) ∈ ℝ^*Q*^ is the time-series of the *j*-th source amplitude of sample *i* and ***n**_i_*(*t*) ∈ ∝^*P*^ is the contamination due to noise. Denoting 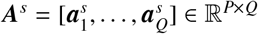, this model is conveniently written in matrix form:

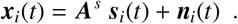

Since we assume that ℝ^*P*^ is the co-product of the source and noise subspaces (they are not ‘mixed’) and that the noise subspace is the same for each sample, the noise writes 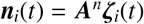 with 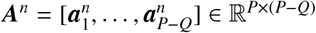 and 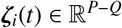. Denoting 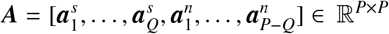 and 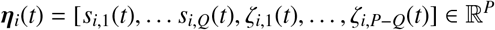, the generative model can be rewritten as:

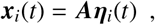

The matrix ***A*** is invertible since the source and noise subspaces span all ℝ^*P*^.

### 5.2. Consistency of SPoC regression model

#### Statement

We consider the previous generative model ***x**_i_*(*t*) = ***Aη**_i_*(*t*) and the outcomes *y*_1_,…, *y_N_*. Define the weighted average 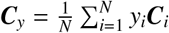 and the average 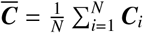. SPoC finds a matrix *W* ∈ ℝ^*P×P*^ solution of the generalized eigenvalue problem:

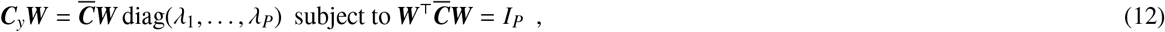

where *λ*_1_,…, *λ_P_* are the generalized eigenvalues. We assume that the eigenvalues are all distinct, and therefore without loss of generality *λ*_1_ > … > *λ_P_*. Under this condition, the generalized eigenvalue problem has a unique solution **W**, and *Q* rows of **W**^*τ*^ match the first *Q* rows of **A**^-1^ up to scale. In particular, the transform **W**^⊤^***x**_i_* recovers the *Q* sources *s_i_*.

*Proof.* We recall the definition 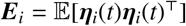, which is block-diagonal with the sources powers *p_i,j_* as coefficient (*j, j*) when *j* ≤ *Q*. We have ***C**_i_* = ***A E**_i_ **A***^⊤^, and therefore 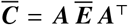 and ***C**_y_* = ***A E**_*y*_ **A***^⊤^, with 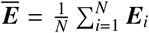 and 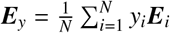 sharing the same block-diagonal structure than the ***E**_i_*. Their lower (*P _ Q*) × (*P _ Q*) diagonal blocks, respectively 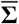 and **Σ**_*y*_, are symmetric matrices. Further, 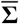 is definite positive, as a linear combination with positive coefficients of definite positive matrices. Hence, 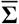 and **Σ**_*y*_ are co-diagonalizable *i.e.* there exists an invertible matrix ***Z*** such that **Σ**_*y*_ = ***ZD_y_Z***^⊤^ and 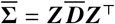. By denoting 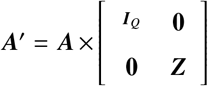, we have that 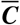 and ***C**_y_* are co-diagonalized by ***A***’. Let ***D*** the diagonal matrix such that 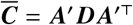. The matrix ***W*** = ***A***’^-τ^***D***^-1/2^ is solution of the generalized eigenvalue problem. By the unicity assumption, SPoC recovers **W** up to a permutation of its columns. The first *Q* rows of ***W***^⊤^ are the first *Q* columns of ***A***’^-τ^, hence the first *Q* rows of ***A***^-1^ up to scale: SPoC recovers the sources.

### 5.3. Consistency of Riemann regression model

#### Statement

We consider the previous generative model ***x**_i_*(*t*) = ***Aη**_i_*(*t*) and the outcomes *y*_1_,…, *y_N_*, which follow a linear model with respect to the log of the source powers: *y_i_* = *β*^⊤^ log(***p**_i_*). Denote the geometric mean 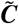 (defined in section 5.5). Let ***v_i_*** the projection of ***C***_2_ on the tangent plane at 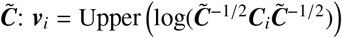. Then, *y_i_* is a linear combination of the coefficients of ***v**_i_*.

*Proof.* First, we note that by invariance, 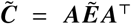, where 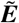 has the same block diagonal structure as the ***E**_i_*’s, and 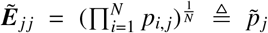 for *j* ≤ *Q*. The vectorization is 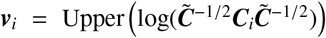. We observe that 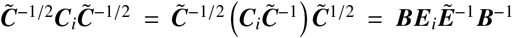 with 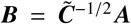. Therefore, 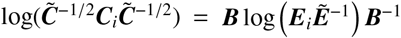, since matrix logarithm is equivariant by similarity. The *Q* values on the diagonal part of 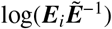 are the log 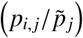. In particular, by denoting 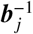 the *j*-th row of ***B***^-1^ and ***b**_j_* the *j*-th column of **B**, we find:

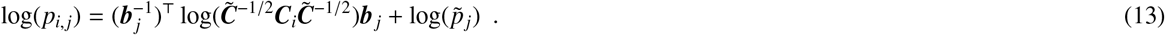

This equation means that log(*p_i,j_*) is obtained as a linear combination of the coefficients in 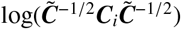, i.e. the coefficients of the vectorization ***v**_i_*. Since *y_i_* is itself a linear combination of the log(*p_i,j_*, the advertised result holds.

As a side note, we have that 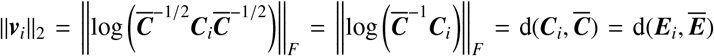, by affine-invariance of the geometric distance *d*(·) (see Appendix 5.5): the norm of ***v**_i_* does not depend on **A**, but only on the log source powers and noise.

### 5.4. Riemannian manifolds

In this work, we consider differentiable manifolds 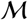 in ℝ^*P*^ of dimension *K*. Intuitively differentiable manifolds are “curved” spaces that locally resemble a flat vector space at each point (see Absil et al. 2009, chap. 3 and Pennec et al. 2006). Examples of differentiable manifolds in ℝ^3^ are curves (one-dimensional manifolds which locally look like a straight line) and surfaces (two-dimensional manifolds which locally look like a plane). More precisely, each point of the manifold 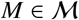 is associated to a vector space called *tangent space* at *M*, denoted 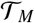. It is the set of derivatives of curves on the manifold passing through *M*. The dimension of 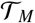 is the dimension of 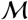, *K*. The differentiable manifold becomes *Riemannian* when each tangent space 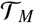 is endowed with a metric, i.e. an inner product 〈·, ·〉_*M*_, giving it an Euclidean structure. This metric is supposed smooth across points on the manifold. We can then define:

- A norm on the tangent space 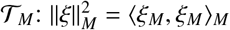 for 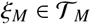.
- The length of a path between two points 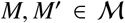: for a path 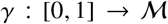 such that *γ*(0) = *M* and *γ*(1) = *M*’, the length of *γ* is 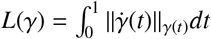. This generalizes the usual notion of path length in Euclidean spaces, where the metric is constant.
- A distance on the manifold 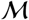, defined as the minimum length of paths: *d*(*M, M*’) = min *L*(*γ*) such that *γ*(0) = *M*, and *γ*(1) = *M*’. This distance is called the *geodesic* distance. If 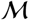 is an Euclidean space, this distance is simply the usual Euclidean distance: *d*(*M, M*’) = ||*M - M*’||_2_, achieved when *γ* is a straight line between *M* and *M*’.
- The Frechet mean 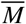 of a set of points 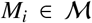 is defined as 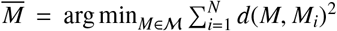. This is a generalization of averaging on manifolds. Indeed, in an Euclidean space, the average 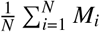 is the Frechet mean of *M*_1_,… *M_N_* with respect to the Euclidean distance *d*(*M, M*’) = ||*M - M*’||_2_. Another example is the geometric mean between positive numbers *a*_1_,…, *a_N_* > 0, given by 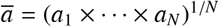, which if the Frechet mean of (*a*_1_,…, *a_N_*) with respect to the distance 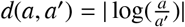.
- The exponential mapping 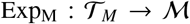 is the operation that maps the tangent space, which has a simple Euclidean structure, to the manifold which might have a much more complicated structure. It satisfies 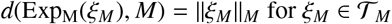 small enough.
- The logarithm mapping 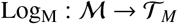 is defined as the reciprocal of the exponential mapping which hence verifies ||Log_M_(*M*’)||_*M*_ = *d*(*M, M*’) for 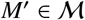 close enough from *M*. It maps the manifold to the tangent space, while preserving the local properties of the manifold.

The logarithm mapping is of crucial importance in practical applications, since it allows to manipulate and store vectors (belonging to the tangent space) instead of points on the manifold. To be more concrete, since each tangent space is a *K*-dimensional Euclidean space, there exists a linear and invertible mapping 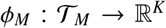 such that 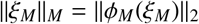 for 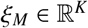. Combining *ϕ_M_* and Log_M_ gives the vectorization operator 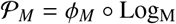 which maps 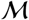 to ℝ^*K*^, and verifies: 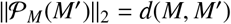 for 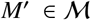. This operator explicitly captures the local Euclidean properties of the Riemannian manifold.

As a final note, all the notions developed above are based on the metric 〈·, 〈〉_*M*_. Different metrics lead to different geodesic distances, Frechet means, exponential and logarithm mapping and vectorization operator. Choosing the right metric for a particular problem may lead to substantial benefits.

### 5.5. The positive definite manifold

In this work, we are interested in one manifold in particular: the manifold of positive definite matrices 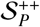 (Förstner and Moonen, 2003). This is not a vector space, as for example the difference of two positive definite matrices may not be positive definite. It is a differentiable manifold of dimension 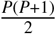, with fixed tangent spaces 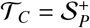 for all 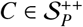. We endow the manifold with the *geometric* metric given by: 〈*P, Q*〉_*C*_ = Tr(*PC*^-1^ *QC*^-1^). This metric leads to closed-form formulas for most Riemannian notions seen above:

- The associated norm generalizes the Froebenius norm: ||*P*||_*I*_ (identity) = ||*P*||_*F*_ (Frobenius) for 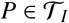.
- The geodesic distance (also called geometric distance) is 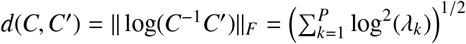, where *λ_k_* are the eigenvalues of *C*^-1^*C*’. This distance is *affine invariant*, i.e. for *W* invertible, *d*(*WCW*^⊤^, *WC*’*W*^⊤^) = *d*(*C, C*’). This is an important property for our purpose: assume that *C* and *C*’ are covariances of some signals *x* and *x*’ ∈ ℝ^*P*^. Affine invariance implies that the distance computed with *x* and *x*’ is the same as the distance computed with any linear transform of the signals *W_x_* and *W_x_*’: the distance is blind to global mixing effects. We also see that the singular matrices act as a barrier for this distance: if *C* or *C*’ is close from being singular, one eigenvalue *λ_k_* goes either to 0 or +∞, and *d*(*C, C*’) goes to infinity.
- For *P* = 1, the Frechet mean is the geometric mean between positive scalars. In higher dimension, no closed-form formula for the Frechet mean has been discovered, but iterative algorithms to compute it are available (Congedo et al., 2016; Jeuris et al., 2012). The mean is also affine invariant, in the sense that 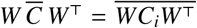. As a consequence, if the matrices *C_i_* were jointly diagonalizable, i.e. *C_i_* = *A*Λ_*i*_*A*^⊤^ with *A* invertible and Λ_*i*_ diagonal, we would have 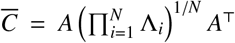. This property is used in the proof of consistency of the Riemann regression model in Appendix 5.3.
- The logarithm mapping is given by 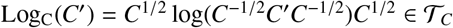, and the vectorization operator is 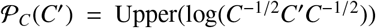, where Upper(*M*) is the vector of size 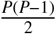 containing the upper triangular coefficients of *M*, with off-diagonal terms weighted by a factor 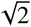. Once again, if *C* and *C*’ are covariances of *x* and *x*’, it amounts to whitening *x*’ with *C*, and then applying a “spectral” non-linear transform on the resulting covariance, where the transform only changes the eigenvalues and not the eigenvectors.

## Footnotes

1 https://github.com/DavidSabbagh/meeg_power_regression.git

## References

Absil, P.-A., Mahony, R., and Sepulchre, R. (2009). Optimization algorithms on matrix manifolds. Princeton University Press.

Akam, T. and Kullmann, D. M. (2014). Oscillatory multiplexing of population codes for selective communication in the mammalian brain. Nature Reviews Neuroscience, 15(2):111.

Allaire, J., Ushey, K., and Tang, Y. (2019). reticulate: Interface to ‘Python’. R package version 1.11.

Andersen, L. M., Pedersen, M. N., Sandberg, K., and Overgaard, M. (2015). Occipital meg activity in the early time range (< 300 ms) predicts graded changes in perceptual consciousness. Cerebral Cortex, 26(6):2677–2688.

Attia, Z. I., Friedman, P. A., Noseworthy, P. A., Lopez-Jimenez, F., Ladewig, D. J., Satam, G., Pellikka, P. A., Munger, T. M., Asirvatham, S. J., Scott, C. G., Carter, R. E., and Kapa, S. (2019). Age and sex estimation using artificial intelligence from standard 12-lead ecgs. Circulation: Arrhythmia and Electrophysiology, 12(9):e007284.

Baillet, S. (2017). Magnetoencephalography for brain electrophysiology and imaging. Nature Neuroscience, 20:327 EP

Barachant, A., Bonnet, S., Congedo, M., and Jutten, C. (2011). Multiclass brain-computer interface classification by riemannian geometry. IEEE Transactions on Biomedical Engineering, 59(4):920–928.

Barachant, A., Bonnet, S., Congedo, M., and Jutten, C. (2013). Classification of covariance matrices using a riemannian-based kernel for bci applications. Neurocomputing, 112:172–178.

Belouchrani, A., Abed-Meraim, K., Cardoso, J.-F., and Moulines, E. (1997). A blind source separation technique using second-order statistics. IEEE Transactions on signal processing, 45(2):434–444.

Besserve, M., Jerbi, K., Laurent, F., Baillet, S., Martinerie, J., and Garnero, L. (2007). Classification methods for ongoing eeg and meg signals. Biological research, 40(4):415–437.

Blankertz, B., Tomioka, R., Lemm, S., Kawanabe, M., and Muller, K. (2008). Optimizing spatial filters for robust eeg single-trial analysis. IEEE Signal Processing Magazine, 25(1):41–56.

Brookes, M. J., Woolrich, M., Luckhoo, H., Price, D., Hale, J. R., Stephenson, M. C., Barnes, G. R., Smith, S. M., and Morris, P. G. (2011). Investigating the electrophysiological basis of resting state networks using magnetoencephalography. Proceedings of the National Academy of Sciences, 108(40):16783–16788.

Buzsáki, G. and Draguhn, A. (2004). Neuronal oscillations in cortical networks. science, 304(5679):1926–1929.

Buzsáki, G. and Mizuseki, K. (2014). The log-dynamic brain: how skewed distributions affect network operations. Nature Reviews Neuroscience, 15(4):264.

Buzsáki, G. and Watson, B. O. (2012). Brain rhythms and neural syntax: implications for efficient coding of cognitive content and neuropsychiatric disease. Dialogues in clinical neuroscience, 14(4):345.

Bzdok, D., Engemann, D., Grisel, O., Varoquaux, G., and Thirion, B. (2018). Prediction and inference diverge in biomedicine: Simulations and real-world data.

Bzdok, D. and Ioannidis, J. P. (2019). Exploration, inference, and prediction in neuroscience and biomedicine. Trends in neurosciences.

Chen, Y., Wiesel, A., Eldar, Y. C., and Hero, A. O. (2010). Shrinkage algorithms for mmse covariance estimation. IEEE Transactions on Signal Processing, 58(10):5016–5029.

Cichy, R. M., Ramirez, F. M., and Pantazis, D. (2015). Can visual information encoded in cortical columns be decoded from magnetoencephalography data in humans? Neuroimage, 121:193–204.

Clarke, E. and Sherrill-Mix, S. (2017). ggbeeswarm: Categorical Scatter (Violin Point) Plots. R package version 0.6.0.

Coles, M. G. and Rugg, M. D. (1995). Event-related brain potentials: An introduction. Oxford University Press.

Congedo, M., Barachant, A., and Andreev, A. (2013). A new generation of brain-computer interface based on Riemannian geometry. arXiv e-prints.

Congedo, M., Barachant, A., and Bhatia, R. (2017). Riemannian geometry for EEG-based brain-computer interfaces; a primer and a review. Brain-Computer Interfaces, 4(3):155–174.

Congedo, M., Phlypo, R., and Barachant, A. (2016). A fixed-point algorithm for estimating power means of positive definite matrices. In 2016 24th European Signal Processing Conference (EUSIPCO), pages 2106–2110.

da Silva, F. L. (2013). Eeg and meg: relevance to neuroscience. Neuron, 80(5):1112–1128.

Dadi, K., Rahim, M., Abraham, A., Chyzhyk, D., Milham, M., Thirion, B., Varoquaux, G., Initiative, A. D. N., et al. (2019). Benchmarking functional connectome-based predictive models for resting-state fmri. NeuroImage, 192:115–134.

Dähne, S., Bießmann, F., Meinecke, F. C., Mehnert, J., Fazli, S., and Müller, K.-R. (2013). Integration of multivariate data streams with bandpower signals. IEEE Transactions on Multimedia, 15(5):1001–1013.

Dähne, S., Meinecke, F. C., Haufe, S., Höhne, J., Tangermann, M., Müller, K.-R., and Nikulin, V. V. (2014a). Spoc: a novel framework for relating the amplitude of neuronal oscillations to behaviorally relevant parameters. Neuroimage, 86:111–122.

Dähne, S., Nikulin, V. V., Ramírez, D., Schreier, P. J., Müller, K.-R., and Haufe, S. (2014b). Finding brain oscillations with power dependencies in neuroimaging data. NeuroImage, 96:334–348.

de Cheveigné, A. and Parra, L. C. (2014). Joint decorrelation, a versatile tool for multichannel data analysis. Neuroimage, 98:487–505.

Dehghani, N., Bédard, C., Cash, S. S., Halgren, E., and Destexhe, A. (2010). Comparative power spectral analysis of simultaneous elecroencephalo-graphic and magnetoencephalographic recordings in humans suggests non-resistive extracellular media. Journal of computational neuroscience, 29(3):405–421.

Delorme, A., Palmer, J., Onton, J., Oostenveld, R., and Makeig, S. (2012). Independent eeg sources are dipolar. PloS one, 7(2):e30135.

Dmochowski, J. P., Sajda, P., Dias, J., and Parra, L. C. (2012). Correlated components of ongoing eeg point to emotionally laden attention-a possible marker of engagement? Frontiers in human neuroscience, 6:112.

Engemann, D. A. and Gramfort, A. (2015). Automated model selection in covariance estimation and spatial whitening of meg and eeg signals. Neuroimage, 108:328–342.

Engemann, D. A., Raimondo, F., King, J.-R., Rohaut, B., Louppe, G., Faugeras, F., Annen, J., Cassol, H., Gosseries, O., Fernandez-Slezak, D., Laureys, S., Naccache, L., Dehaene, S., and Sitt, J. D. (2018). Robust EEG-based cross-site and cross-protocol classification of states of consciousness. Brain, 141(11):3179–3192.

Fischl, B. (2012). FreeSurfer. Neuroimage, 62(2):774–781.

Förstner, W. and Moonen, B. (2003). A metric for covariance matrices. In Geodesy-The Challenge of the 3rd Millennium, pages 299–309. Springer.

Fruehwirt, W., Gerstgrasser, M., Zhang, P., Weydemann, L., Waser, M., Schmidt, R., Benke, T., Dal-Bianco, P., Ransmayr, G., Grossegger, D., et al. (2017). Riemannian tangent space mapping and elastic net regularization for cost-effective eeg markers of brain atrophy in alzheimer’s disease. arXiv preprint arXiv:1711.08359.

Fukunaga, K. (1990). Chapter 2 -random vectors and their properties. In Fukunaga, K., editor, Introduction to Statistical Pattern Recognition (Second Edition), pages 11–50. Academic Press, Boston, second edition edition.

Garcés, P., López-Sanz, D., Maestú, F., and Pereda, E. (2017). Choice of magnetometers and gradiometers after signal space separation. Sensors, 17(12):2926.

Gelman, A. et al. (2005). Analysis of variance — why it is more important than ever. The annals of statistics, 33(1):1–53.

Golub, G. H., Heath, M., and Wahba, G. (1979). Generalized cross-validation as a method for choosing a good ridge parameter. Technometrics, 21(2):215–223.

Gramfort, A., Luessi, M., Larson, E., Engemann, D. A., Strohmeier, D., Brodbeck, C., Parkkonen, L., and Hämäläinen, M. S. (2014). MNE software for processing MEG and EEG data. NeuroImage, 86:446–460.

Gross, J., Baillet, S., Barnes, G. R., Henson, R. N., Hillebrand, A., Jensen, O., Jerbi, K., Litvak, V., Maess, B., Oostenveld, R., et al. (2013). Good practice for conducting and reporting meg research. Neuroimage, 65:349–363.

Grosse-Wentrup‪, M. and Buss, M. (2008). Multiclass common spatial patterns and information theoretic feature extraction. IEEE Transactions on Biomedical Engineering, 55(8):1991–2000.

Halme, H.-L. and Parkkonen, L. (2018). Across-subject offline decoding of motor imagery from meg and eeg. Scientific reports, 8(1):1–12.

Hämäläinen, M., Hari, R., Ilmoniemi, R. J., Knuutila, J., and Lounasmaa, O. V. (1993). Magnetoencephalography—theory, instrumentation, and applications to noninvasive studies of the working human brain. Reviews of modern Physics, 65(2):413.

Hämäläinen, M. and Ilmoniemi, R. (1984). Interpreting magnetic fields of the brain: minimum norm estimates. Technical Report TKK-F-A559, Helsinki University of Technology.

Hämäläinen, M. S. and Ilmoniemi, R. J. (1994). Interpreting magnetic fields of the brain: minimum norm estimates. Medical & biological engineering & computing, 32(1):35–42.

Harati, A., Lopez, S., Obeid, I., Picone, J., Jacobson, M., and Tobochnik, S. (2014). The tuh eeg corpus: A big data resource for automated eeg interpretation. In 2014 IEEE Signal Processing in Medicine and Biology Symposium (SPMB), pages 1–5. IEEE.

Hastie, T., Tibshirani, R., Friedman, J., and Franklin, J. (2005). The elements of statistical learning: data mining, inference and prediction. The Mathematical Intelligencer, 27(2):83–85.

Haufe, S., Dähne, S., and Nikulin, V. V. (2014a). Dimensionality reduction for the analysis of brain oscillations. NeuroImage, 101:583–597.

Haufe, S., Meinecke, F., Görgen, K., Dähne, S., Haynes, J.-D., Blankertz, B., and Bießmann, F. (2014b). On the interpretation of weight vectors of linear models in multivariate neuroimaging. NeuroImage, 87:96–110.

Hauk, O. and Stenroos, M. (2014). A framework for the design of flexible cross-talk functions for spatial filtering of eeg/meg data: Deflect. Human brain mapping, 35(4):1642–1653.

He, B. J., Zempel, J. M., Snyder, A. Z., and Raichle, M. E. (2010). The temporal structures and functional significance of scale-free brain activity. Neuron, 66(3):353–369.

He, T., Kong, R., Holmes, A. J., Nguyen, M., Sabuncu, M. R., Eickhoff, S. B., Bzdok, D., Feng, J., and Yeo, B. T. (2019). Deep neural networks and kernel regression achieve comparable accuracies for functional connectivity prediction of behavior and demographics. NeuroImage, page 116276.

Hild II, K. E. and Nagarajan, S. S. (2009). Source localization of eeg/meg data by correlating columns of ica and lead field matrices. IEEE Transactions on Biomedical Engineering, 56(11):2619–2626.

Hipp, J. F., Hawellek, D. J., Corbetta, M., Siegel, M., and Engel, A. K. (2012). Large-scale cortical correlation structure of spontaneous oscillatory activity. Nature neuroscience, 15(6):884.

Hoerl, A. E. and Kennard, R. W. (1970). Ridge regression: Biased estimation for nonorthogonal problems. Technometrics, 12(1):55–67.

Honey, C. J., Kötter, R., Breakspear, M., and Sporns, O. (2007). Network structure of cerebral cortex shapes functional connectivity on multiple time scales. Proceedings of the National Academy of Sciences, 104(24):10240–10245.

Hotelling, H. (1992). Relations between two sets of variates. In Breakthroughs in statistics, pages 162–190. Springer.

Hyvärinen, A., Karhunen, J., and Oja, E. (2004). Independent component analysis, volume 46. John Wiley & Sons.

Hyvärinen, A. and Oja, E. (2000). Independent component analysis: algorithms and applications. Neural networks, 13(4-5):411–430.

Jas, M., Engemann, D. A., Bekhti, Y., Raimondo, F., and Gramfort, A. (2017). Autoreject: Automated artifact rejection for MEG and EEG data. NeuroImage, 159:417–429.

Jas, M., Larson, E., Engemann, D. A., Leppäkangas, J., Taulu, S., Hämäläinen, M., and Gramfort, A. (2018). A reproducible MEG/EEG group study with the MNE software: Recommendations, quality assessments, and good practices. Frontiers in Neuroscience, 12(AUG):1–18.

Jeuris, B., Vandebril, R., and Vandereycken, B. (2012). A survey and comparison of contemporary algorithms for computing the matrix geometric mean. Electronic Transactions on Numerical Analysis, 39(ARTICLE):379–402.

Khan, S., Hashmi, J. A., Mamashli, F., Michmizos, K., Kitzbichler, M. G., Bharadwaj, H., Bekhti, Y., Ganesan, S., Garel, K.-L. A., Whitfield-Gabrieli, S., et al. (2018). Maturation trajectories of cortical resting-state networks depend on the mediating frequency band. NeuroImage, 174:57–68.

Kietzmann, T. C., Spoerer, C. J., Sörensen, L. K. A., Cichy, R. M., Hauk, O., and Kriegeskorte, N. (2019). Recurrence is required to capture the representational dynamics of the human visual system. Proceedings of the National Academy of Sciences.

King, J.-R. and Dehaene, S. (2014). Characterizing the dynamics of mental representations: the temporal generalization method. Trends in cognitive sciences, 18(4):203–210.

King, J.-R., Faugeras, F., Gramfort, A., Schurger, A., El Karoui, I., Sitt, J., Rohaut, B., Wacongne, C., Labyt, E., Bekinschtein, T., et al. (2013). Single-trial decoding of auditory novelty responses facilitates the detection of residual consciousness. Neuroimage, 83:726–738.

King, J.-R., Gwilliams, L., Holdgraf, C., Sassenhagen, J., Barachant, A., Engemann, D., Larson, E., and Gramfort, A. (2018). Encoding and decoding neuronal dynamics: Methodological framework to uncover the algorithms of cognition.

Koles, Z. J. (1991). The quantitative extraction and topographic mapping of the abnormal components in the clinical eeg. Electroencephalography and clinical Neurophysiology, 79(6):440–447.

Larson-Prior, L. J., Oostenveld, R., Della Penna, S., Michalareas, G., Prior, F., Babajani-Feremi, A., Schoffelen, J.-M., Marzetti, L., de Pasquale, F., Di Pompeo, F., et al. (2013). Adding dynamics to the Human Connectome Project with MEG. Neuroimage, 80:190–201.

Liem, F., Varoquaux, G., Kynast, J., Beyer, F., Masouleh, S. K., Huntenburg, J. M., Lampe, L., Rahim, M., Abraham, A., Craddock, R. C., Riedel-Heller, S., Luck, T., Loeffler, M., Schroeter, M. L., Witte, A. V., Villringer, A., and Margulies, D. S. (2017). Predicting brain-age from multimodal imaging data captures cognitive impairment. NeuroImage, 148:179–188.

Lin, F.-H., Belliveau, J. W., Dale, A. M., and Hämäläinen, M. S. (2006). Distributed current estimates using cortical orientation constraints. Human brain mapping, 27(1):1–13.

Linkenkaer-Hansen, K., Nikouline, V. V., Palva, J. M., and Ilmoniemi, R. J. (2001). Long-range temporal correlations and scaling behavior in human brain oscillations. Journal of Neuroscience, 21(4):1370–1377.

Lotte, F., Bougrain, L., Cichocki, A., Clerc, M., Congedo, M., Rakotomamonjy, A., and Yger, F. (2018). A review of classification algorithms for EEG-based brain-computer interfaces: a 10 year update. Journal of Neural Engineering, 15(3):031005.

Lotte, F., Congedo, M., Lécuyer, A., Lamarche, F., and Arnaldi, B. (2007). A review of classification algorithms for EEG-based brain-computer interfaces. Journal of Neural Engineering, 4(2):R1–R13.

Makeig, S., Bell, A. J., Jung, T.-P., and Sejnowski, T. J. (1995). Independent component analysis of electroencephalographic data. In Proceedings of the 8th International Conference on Neural Information Processing Systems, NIPS’95, pages 145–151, Cambridge, MA, USA. MIT Press.

Makeig, S., Bell, A. J., Jung, T.-P., and Sejnowski, T. J. (1996). Independent component analysis of electroencephalographic data. In Advances in neural information processing systems, pages 145–151.

Makeig, S., Jung, T.-P., Bell, A. J., Ghahremani, D., and Sejnowski, T. J. (1997). Blind separation of auditory event-related brain responses into independent components. Proceedings of the National Academy of Sciences, 94(20):10979–10984.

Mazaheri, A., Segaert, K., Olichney, J., Yang, J.-C., Niu, Y.-Q., Shapiro, K., and Bowman, H. (2018). Eeg oscillations during word processing predict mci conversion to alzheimer’s disease. NeuroImage: Clinical, 17:188–197.

Mosher, J. C., Leahy, R. M., and Lewis, P. S. (1999). EEG and MEG: forward solutions for inverse methods. IEEE Transactions on Biomedical Engineering, 46(3):245–259.

Mosher, J. C., Lewis, P. S., and Leahy, R. M. (1992). Multiple dipole modeling and localization from spatio-temporal MEG data. IEEE Transactions on Biomedical Engineering, 39(6):541–557.

Näätänen, R. (1975). Selective attention and evoked potentials inhumans—a critical review. Biological Psychology, 2(4):237–307.

Nikulin, V. V., Nolte, G., and Curio, G. (2011). A novel method for reliable and fast extraction of neuronal eeg/meg oscillations on the basis of spatio-spectral decomposition. NeuroImage, 55(4):1528–1535.

Nolte, G., Ziehe, A., Meinecke, F., and Müller, K.-R. (2006). Analyzing coupled brain sources: Distinguishing true from spurious interaction. In Weiss, Y., Schölkopf, B., and Platt, J. C., editors, Advances in Neural Information Processing Systems 18, pages 1027–1034. MIT Press.

Olivetti, E., Kia, S. M., and Avesani, P. (2014). Meg decoding across subjects. In 2014 International Workshop on Pattern Recognition in Neuroimaging, pages 1–4. IEEE.

Oostenveld, R., Fries, P., Maris, E., and Schoffelen, J.-M. (2011). Fieldtrip: open source software for advanced analysis of meg, eeg, and invasive electrophysiological data. Computational intelligence and neuroscience, 2011:1.

Panzeri, S., Brunel, N., Logothetis, N. K., and Kayser, C. (2010). Sensory neural codes using multiplexed temporal scales. Trends in neurosciences, 33(3):111–120.

Parra, L. C., Spence, C. D., Gerson, A. D., and Sajda, P. (2005). Recipes for the linear analysis of eeg. Neuroimage, 28(2):326–341.

Pedregosa, F., Varoquaux, G., Gramfort, A., Michel, V., Thirion, B., Grisel, O., Blondel, M., Prettenhofer, P., Weiss, R., Dubourg, V., Vanderplas, J., Passos, A., Cournapeau, D., Brucher, M., Perrot, M., and Duchesnay, E. (2011). Scikit-learn: Machine learning in Python. Journal of Machine Learning Research, 12:2825–2830.

Pennec, X., Fillard, P., and Ayache, N. (2006). A riemannian framework for tensor computing. International Journal of computer vision, 66(1):41–66.

Pernet, C. R., Garrido, M., Gramfort, A., Maurits, N., Michel, C., Pang, E., Salmelin, R., Schoffelen, J. M., Valdes-Sosa, P. A., and Puce, A. (2018). Best Practices in Data Analysis and Sharing in Neuroimaging using MEEG.

Polich, J. and Kok, A. (1995). Cognitive and biological determinants of p300: an integrative review. Biological psychology, 41(2):103–146.

R Core Team (2019). R: A Language and Environment for Statistical Computing. R Foundation for Statistical Computing, Vienna, Austria.

Roberts, J. A., Boonstra, T W., and Breakspear, M. (2015). The heavy tail of the human brain. Current opinion in neurobiology, 31:164–172.

Rodrigues, P. L. C., Bouchard, F., Congedo, M., and Jutten, C. (2017). Dimensionality Reduction for BCI classification using Riemannian geometry. In 7th Graz Brain-Computer Interface Conference (BCI 2017), Graz, Austria. Gernot R. Müller&-Putz.

Rodrigues, P. L. C., Congedo, M., and Jutten, C. (2018). Multivariate time-series analysis via manifold learning. In 2018 IEEE Statistical Signal Processing Workshop (SSP), pages 573–577. IEEE.

Rodrigues, P. L. C., Jutten, C., and Congedo, M. (2019). Riemannian procrustes analysis: Transfer learning for brain-computer interfaces. IEEE Transactions on Biomedical Engineering, 66(8):2390–2401.

Roy, Y., Banville, H., Albuquerque, I., Gramfort, A., Falk, T. H., and Faubert, J. (2019). Deep learning-based electroencephalography analysis: a systematic review. Journal of Neural Engineering, 16(5):051001.

Sabbagh, D., Ablin, P., Varoquaux, G., Gramfort, A., and Engemann, D. (2019). Manifold-regression to predict from meg/eeg brain signals without source modeling. arXiv Preprint 1906.02687v2.

Sami, S., Williams, N., Hughes, L. E., Cope, T. E., Rittman, T., Coyle-Gilchrist, I. T., Henson, R. N., and Rowe, J. B. (2018). Neurophysiological signatures of alzheimer’s disease and frontotemporal lobar degeneration: pathology versus phenotype. Brain, 141(8):2500–2510.

Schirrmeister, R., Gemein, L., Eggensperger, K., Hutter, F., and Ball, T. (2017). Deep learning with convolutional neural networks for decoding and visualization of eeg pathology. In 2017 IEEE Signal Processing in Medicine and Biology Symposium (SPMB), pages 1–7. IEEE.

Schoffelen, J.-M., Poort, J., Oostenveld, R., and Fries, P. (2011). Selective movement preparation is subserved by selective increases in corticomuscular gamma-band coherence. Journal of Neuroscience, 31(18):6750–6758.

Schulz, M.-A., Yeo, T., Vogelstein, J., Mourao-Miranada, J., Kather, J., Kording, K., Richards, B. A., and Bzdok, D. (2019). Deep learning for brains?: Different linear and nonlinear scaling in uk biobank brain images vs. machine-learning datasets. bioRxiv, page 757054.

Shafto, M. A., Tyler, L. K., Dixon, M., Taylor, J. R., Rowe, J. B., Cusack, R., Calder, A. J., Marslen-Wilson, W. D., Duncan, J., Dalgleish, T., et al. (2014). The Cambridge Centre for Ageing and Neuroscience (Cam-CAN) study protocol: a cross-sectional, lifespan, multidisciplinary examination of healthy cognitive ageing. BMC neurology, 14(1):204.

Siegel, M., Donner, T. H., and Engel, A. K. (2012). Spectral fingerprints of large-scale neuronal interactions. Nature Reviews Neuroscience, 13(2):121.

Stewart, A. X., Nuthmann, A., and Sanguinetti, G. (2014). Single-trial classification of eeg in a visual object task using ica and machine learning. Journal of neuroscience methods, 228:1–14.

Stokes, M. G., Wolff, M. J., and Spaak, E. (2015). Decoding rich spatial information with high temporal resolution. Trends in cognitive sciences, 19(11):636–638.

Strubell, E., Ganesh, A., and McCallum, A. (2019). Energy and policy considerations for deep learning in nlp. arXivpreprint arXiv:1906.02243.

Subasi, A. and Gursoy, M. I. (2010). Eeg signal classification using pca, ica, lda and support vector machines. Expert systems with applications, 37(12):8659–8666.

Tallon-Baudry, C. and Bertrand, O. (1999). Oscillatory gamma activity in humans and its role in object representation. Trends in cognitive sciences, 3(4):151–162.

Tangermann, M. W., Krauledat, M., Grzeska, K., Sagebaum, M., Vidaurre, C., Blankertz, B., and Müller, K.-R. (2008). Playing pinball with non-invasive bci. In Proceedings of the 21st International Conference on Neural Information Processing Systems, pages 1641–1648. Citeseer.

Taulu, S. and Kajola, M. (2005). Presentation of electromagnetic multichannel data: the signal space separation method. Journal of Applied Physics, 97(12):124905.

Taylor, J. R., Williams, N., Cusack, R., Auer, T., Shafto, M. A., Dixon, M., Tyler, L. K., Henson, R. N., et al. (2017). The Cambridge Centre for Ageing and Neuroscience (Cam-CAN) data repository: structural and functional MRI, MEG, and cognitive data from a cross-sectional adult lifespan sample. Neuroimage, 144:262–269.

Thavikulwat, A. T., Lopez, P., Caruso, R. C., and Jeffrey, B. G. (2015). The effects of gender and age on the range of the normal human electro-oculogram. Documenta Ophthalmologica, 131(3):177–188.

Tipping, M. E. and Bishop, C. M. (1999). Probabilistic principal component analysis. Journal of the Royal Statistical Society: Series B (Statistical Methodology), 61(3):611–622.

Uusitalo, M. A. and Ilmoniemi, R. J. (1997). Signal-space projection method for separating MEG or EEG into components. Medical and Biological Engineering and Computing, 35(2):135–140.

Van Veen, B. D. and Buckley, K. M. (1988). Beamforming: A versatile approach to spatial filtering. IEEE assp magazine, 5(2):4–24.

van Vliet, M. and Salmelin, R. (2019). Post-hoc modification of linear models: Combining machine learning with domain information to make solid inferences from noisy data. NeuroImage, page 116221.

van Wassenhove, V. (2016). Temporal cognition and neural oscillations. Current Opinion in Behavioral Sciences, 8:124–130.

Varoquaux, G., Raamana, P. R., Engemann, D. A., Hoyos-Idrobo, A., Schwartz, Y., and Thirion, B. (2017). Assessing and tuning brain decoders: cross-validation, caveats, and guidelines. NeuroImage, 145:166–179.

Wang, Y. and Makeig, S. (2009). Predicting intended movement direction using eeg from human posterior parietal cortex. In International Conference on Foundations of Augmented Cognition, pages 437–446. Springer.

Wardle, S. G., Kriegeskorte, N., Grootswagers, T., Khaligh-Razavi, S.-M., and Carlson, T. A. (2016). Perceptual similarity of visual patterns predicts dynamic neural activation patterns measured with meg. Neuroimage, 132:59–70.

Westner, B. U., Dalal, S. S., Hanslmayr, S., and Staudigl, T. (2018). Across-subjects classification of stimulus modality from human meg high frequency activity. PLoS computational biology, 14(3):e1005938.

Wickham, H. (2016). ggplot2: Elegant Graphics for Data Analysis. Springer-Verlag New York.

Wolpaw, J. R., McFarland, D. J., Neat, G. W., and Forneris, C. A. (1991). An eeg-based brain-computer interface for cursor control. Electroencephalography and clinical neurophysiology, 78(3):252–259.

Woo, C.-W., Chang, L. J., Lindquist, M. A., and Wager, T. D. (2017). Building better biomarkers: brain models in translational neuroimaging. Nature neuroscience, 20(3):365.

Woolrich, M., Hunt, L., Groves, A., and Barnes, G. (2011). Meg beamforming using bayesian pca for adaptive data covariance matrix regularization. Neuroimage, 57(4):1466–1479.

Yger, F., Berar, M., and Lotte, F. (2017). Riemannian approaches in brain-computer interfaces: A review. IEEE Transactions on Neural Systems and Rehabilitation Engineering, 25(10):1753–1762.

